# Thermodynamic Parameters Contributions of Single Internal Mismatches In RNA/DNA Hybrid Duplexes

**DOI:** 10.1101/2022.11.25.517909

**Authors:** Tongjun Xiang, Huibao Feng, Xin-hui Xing, Chong Zhang

## Abstract

Nucleic acid is a fundamental component of life. The thermodynamic is a basic physical quantity for nucleic acid and has been adopted in varieties of biotechnological applications. However, the parameters predicting thermodynamic parameters of RNA/DNA hybrid duplexes is incomplete and thus lead to the inability to proper predicting the properties. To address this problem, we measured for 99 RNA/DNA hybrid duplexes with 11 nucleotide length in a 1 M NaCl buffer using UV absorbance thermal denaturation. Thermodynamic parameters were obtained by fitting absorbance versus temperature profiles using SciPy non-linear curve fit. A dataset of 228 duplexes were constructed combining with literature experimental thermodynamic parameters for fitting a full set of all 113 possible Watson-Crick and single internal mismatch involving nearest-neighbor parameters using singular value decomposition. The new nearest-neighbor parameters predict thermodynamic parameters of RNA/DNA hybrid duplexes within an average error of 0.26 kcal/mol (3.55%) in free energy at 37°C (ΔG°_37_), 5.05 kcal/mol (6.70%) in enthalpy (ΔH°), 16.3 cal/mol·K (7.50%) in entropy (ΔS°) and 1.14°C (2.91%) in melting temperature (T_m_). The highest accuracy of all parameters reported. We observed a trend of increased stability for mismatch involving pairs of rG/dT >> rU/dG > rG/dA > rG/dG ≈ rA/dC ≈ rA/dG > rC/dA ≈ rC/dT > rU/dT ≈ rU/dC >> rA/dA >> rC/dC, which can be explained by different hydrogen bond conditions in mismatch pairs. We further compared the stability with RNA/RNA and DNA/DNA duplexes and found a backbone structure depended pattern in these three kinds of nucleic acid duplexes. This nearest-neighbor parameter set could enhance thermodynamical understanding of RNA/DNA combination and have potential usage in predicting on- and off-target RNA-DNA binding activity when applied to biotechnological applications such as CRISPR/Cas systems, RNAi and qPCR.

## INTRODUCTION

As the foundation of life, nucleic acids play essential roles in various biological progresses (1–4). Engineered nucleic acids are also broadly used as therapeutic agents (5–10), diagnostics reagents (11–16), building blocks for precisely patterned structures (17–21) and complex spatiotemporal circuits (22–26), etc. For either the physiological processes or the practical applications, the thermodynamics of base pairing, which is closely related to the stability of the nucleic acid secondary structures, to a large extent, determines the fundamental macroscopic property of the nucleic acids. Therefore, there is a critical need to obtain thermodynamic parameters of nucleic acids in order to deepen our understanding of life and to guide the rational design of nucleic acid structures.

For the past three decades, nucleic acid thermodynamics has been adequately approximated by the nearest-neighbor model (27–30), which assumes that the thermodynamic property of a given base pair is only affected by its adjacent base pair, hence the corresponding parameters are pairwise additive (31). This model has been extensively tested and is in agreement with experimental data (32). To date, thermodynamic parameters for Watson-Crick pairs of DNA/DNA (33), RNA/RNA (34) and RNA/DNA (35) have been fully determined. These parameters have greatly contributed to the understanding of biological processes (36,37) and are widely used in structure prediction (38,39), stability prediction (40,41), moderate expansion (42), annealing optimization (43), etc. As for non-Watson-Crick pairs, or pairs involving mismatches, which are commonly found in RNA secondary structures (44–46), also play important roles in applications involving nucleic acid-nucleic acid annealing process, such as real-time PCR (47,48), RNA interference (49,50), CRISPR/Cas system (51–53), etc. Although a full set of thermodynamic parameters for RNA/RNA and DNA/DNA mismatches have been reported (54–59), the parameters for RNA/DNA hybrid duplexes are not sufficiently tested (60), with 8 types of mismatches (rA/dC, rA/dG, rU/dC, rU/dG, rC/dA, rC/dT, rG/dA and rG/dT) undetermined, leading to the inability of predicting the properties of hybrid duplexes containing these mismatches. Thus, a comprehensive measurement of nearest-neighbor parameters is urgently required.

To address the above issue, we assayed the melting curves of 99 different 11-bp RNA/DNA hybrid duplexes via UV-VIS spectrophotometer, each with an internal mismatch base pair in the central position. Moreover, we combined our data with previously reported results, including 72 duplexes with only Watson-Crick pairs and 57 duplexes containing mismatches from the literature (35,60), to generate a comprehensive thermodynamic dataset of 228 duplexes. This dataset was further analyzed using singular value decomposition (SVD) (61,62) to obtain a full set of 113 nearest-neighbor parameters of RNA/DNA hybrid duplexes (16 types of Watson-Crick pairs, 96 types of mismatches and the regression initiation thermodynamic parameter). The thermodynamic properties calculated using determined nearest-neighbor parameters were highly consistent with the experimental values, indicating the reliability of our results. Therefore, based on these parameters, we are able to predict the stability of RNA/DNA hybrid duplexes. Furthermore, these parameters have great potential to be applied in multiple situations, such as designing crRNAs with tailored binding activity, optimization of RNA primers and probes in RNAi and RT-PCR.

## MATERIALS AND METHODS

### Synthesis and purification of sequences

The RNA and DNA oligonucleotides are synthesized by Tsingke Biotechnology, Beijing and Azenta Life Sciences, Suzhou, respectively. All oligonucleotides are deblocked and HPLC purified. Mass-spectrometry is performed by the suppliers to certify the sequence of oligonucleotides.

### Design of sequences

Each oligonucleotide used in our study is 11 bp in length with the mismatch in central position. Four or more consecutive identical bases is avoided to ensure a complete complement. G/C content is controlled within 30-70% to maintain the melting temperature (T_m_) of duplexes at 10 μM to be between 30 and 60°C, hence the pre- and post-transition region of melting curve could be long enough for fitting upper and lower baselines, therefore, appropriate curve-fitting could be conducted to deduce accurate thermodynamic parameters. To avoid the impact on thermodynamics of secondary structures (63), each sequence was examined using RNAFold to ensure no formation of undesired hairpins, slipped duplexes, homodimers or other possible secondary structures (64). Finally, at least of 10 duplexes with different 5’- and 3’-adjacent Watson-Crick base pairs for each mismatch type are designed to ensure each type of nearest-neighbor sequence is at least 2-fold over-determined. The total size of experimental group is 99 duplexes (33 RNA oligonucleotides and 99 DNA oligonucleotides, see **Supplementary table S1**).

### Measurement of melting curves

The melting buffer used in our study was 1.0 M NaCl, 10 mM Na_2_HPO_4_ and 0.5 mM Na_2_EDTA, and titrated to pH of 7.0 with a 0.1 M HCl. High salt concentrations can remove the length-dependent counterion-condensation effects which influences the proper determination of nearest-neighbor parameters (60). The concentrations of single-stranded oligonucleotides were determined from the UV absorbance measured under 260 nm using a Thermo Scientific NanoDrop One spectrophotometer using Equation (1).

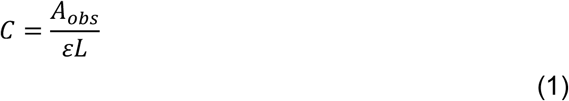

Where A_obs_ is the measured UV absorbance, ε is the extinction coefficients of oligonucleotide, and L is the optical path length.

Extinction coefficients of both RNA and DNA oligonucleotides were predicted based on the nearest-neighbor model (65–67) using the extinction coefficient calculator provided by Integrated DNA Technologies. For each non-self-complementary duplexes, oligonucleotides were mixed at a 1:1 mole ratio with the initial volume of 300 μL. Serial dilutions of this 300 μL sample were made to obtain six different oligonucleotide concentrations over a 20-fold dilution range to make sure UV absorbance reading for each test group was below 2.00 in a 10 mm path length to stay within the linear region of Beer’s Law. Melting curves were measured under 260 nm using a SHIMADZU UV-2700i UV-VIS spectrophotometer with S-1700 Thermoelectric Single Cell Holder in a 10 mm 8 Series Micro Multi-Cells. The melting curves were measured from 10°C to a maximum of 95°C with a heating rate of 1.0°C /min. Read interval was set to 0.5°C for getting enough data for curve-fitting. The total strand concentration (C_T_) of each test group was calculated from the experimental UV absorbance at 85°C, under this temperature we assume the RNA and DNA are both single-stand oligonucleotides, using Equation (1) with the extinction coefficients to be the average extinction coefficients of the two individual strand.

### Determination of thermodynamic parameters for duplexes

Thermodynamic parameters for each duplex were obtained through two different methods: the T_m_^−1^ versus ln C_T_/4 method and Marquardt non-linear least squares curve fit method. The Gibbs free energy (ΔG°), enthalpy (ΔH°) and entropy (ΔS°) can be defined using Van’t Hoff plots by assuming the change in heat capacity (ΔCp°) is zero for the transition equilibrium.

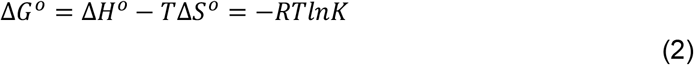

Where R is the gas constant, 1.987 cal/K/mol; T is the temperature; K is the equilibrium constant which can be defined using the ratio of double strands, α, via Equation (3)

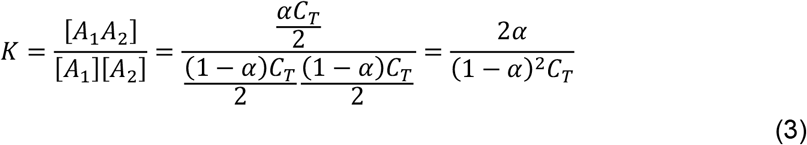

Literatures have indicated that the average ΔCp° for DNA duplexes is approx. 100 cal/mol · K per base pair (68–70). In our study, however, we retained the ΔCp° = 0 assumption because data from UV melting curve are not accurate enough to justify the ΔCp° value. In addition, ΔCp° = 0 assumption is consistent with previous studies (33–35,54–60). Moreover, the ΔCp° = 0 assumption can be demonstrated by the two-state behavior and highly predictive nearest-neighbor parameters determined in this work.

For the T_m_^−1^ versus ln C_T_/4 method, the melting temperature is defined using Tm analysis software by SHIMADZU, it defines the T_m_ as the temperature where melting curve intersect with the average of the upper and lower baselines (71). At this point, the fraction of strands in the native and denatured states is equal to 0.5, where

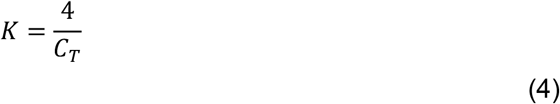

Under this condition, Equation (2) can be rewritten as Equation (5).

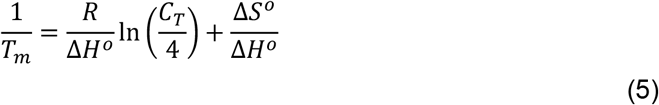

Equation (5) was applied for a linear regression of T_m_^−1^ versus ln C_T_/4 to solve ΔH° and ΔS° for duplexes, the slope was R/ΔH° and the intercept was ΔS°/ΔH°. The Gibbs free energy under 37°C (ΔG°_37_) was the calculated by Equation (6)

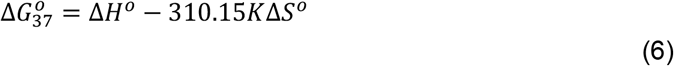

For the Marquardt non-linear least squares method, we assume that the measured UV absorbance of hybrid duplexes is combined by the absorbance of single and double strands which effected by temperature:

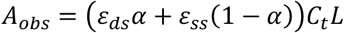

Where A_obs_ is the measured UV absorbance, ε_ds_ and ε_ss_ stand for extinction coefficients of single and double strands, α is the ratio of double strands, and L is the optical path length.

Extinction coefficients of single and double strands are assumed to have linear relations with the temperature. So that we can list:

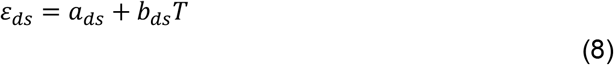

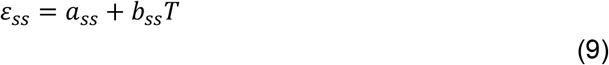

Combining Equation (7, 8 and 9), we can get:

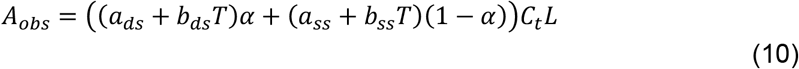

From Equations (2 and 3), we can derive that:

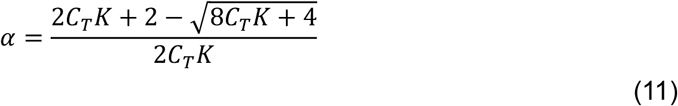

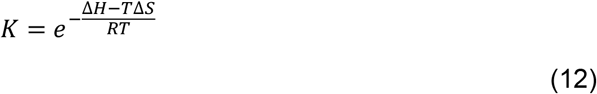

Equations (10, 11 and 12) were used in a non-linear fitting to solve the ΔH° and ΔS° from melting curve.

Thermodynamic parameters from the two methods were checked within a limitation of 15% difference in enthalpy to validate two-state approximation (60,71). Validated data from T_m_^−1^ versus ln C_T_/4 method was used to fit nearest-neighbor parameters. Raw data of the test group in our study for determination of nearest-neighbor contributions of single internal mismatches in RNA/DNA hybrid duplexes are shown in **Supplementary Table S1**.

### Error analysis

Sampling errors of ΔH° and ΔS° for T_m_^−1^ versus ln C_T_/4 plots and curve fits were determined from the linear regression errors using optimization package from SciPy 1.9.1, respectively. Errors of ΔG°_37_ are calculated from errors of ΔH° and ΔS°. Experimental ΔH° and ΔS° parameters are not independently determined, instead, they are highly correlated with a typical R^2^ > 0.99 (71), the errors propagate to ΔG°_37_ are much smaller than would be expected if ΔH° and ΔS° were uncorrelated. The error of can ΔG°_37_ be determined by Equation (13).

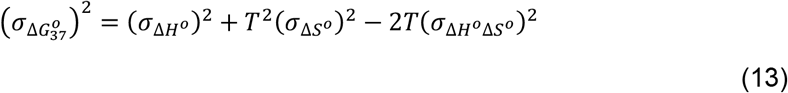

Sampling errors of duplexes from literatures (35,60) were also combined with errors from our experiment to compute the square-root of the sum of the squares for duplex ΔG°_37_, ΔH° and ΔS°, which were the error values used in SVD calculations using MATLAB R2022a.

### Determination of nearest-neighbor parameters

In order to fit a unified set of all 113 types (16 Watson-Crick pairs, 96 mismatch involving pairs and the regression initiation thermodynamic parameter) of nearest-neighbor parameters in RNA/DNA hybrid duplexes, literature reported thermodynamic parameters from Watkins *et al*. (65 groups of Watson-Crick and single internal mismatch containing Duplexes, see **Supplementary Table S2**) and Sugimoto *et al*. (64 groups of Watson-Crick Duplexes, see **Supplementary Table S3**) are combined with our results (**Supplementary Table S1**) to establish a full set of thermodynamic dataset. According to the nearest-neighbor model, thermodynamic parameters for a given sequence can be determined by the summation of duplex initiation and nearest-neighbor interactions between base pairs. For instance, the thermodynamic parameters ΔY°_total_ (where Y = G, H or S) for the duplex sequences (rGAGCAAACGUC/dGACGTCTGCTC) are decomposed as follows:

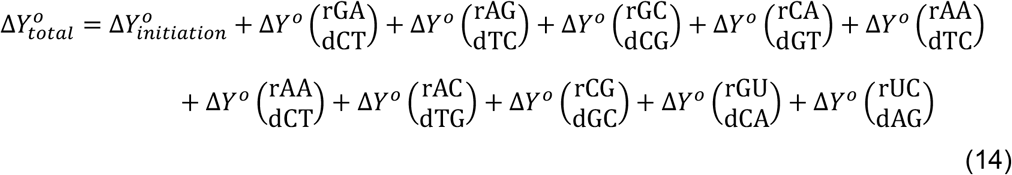

Equation (14) can be constructed into more general Equation (15)

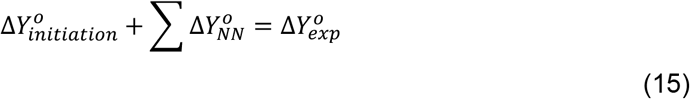

A system of equations based on Equation (15) were used to set up a coefficient matrix as input to MATLAB R2022a. The set of 113 nearest-neighbor parameters (ΔG°_37_, ΔH° and ΔS°) were then solved using SVD. Since none of the sequences have terminal mismatches, each sequence would have two mismatch involving component when calculating thermodynamic parameters. This would make each mismatch type only have seven degrees of freedom instead of eight, resulting the rank deficient of the coefficient matrix (60). Thus, the SVD matrix is reduced to 101 degrees for solving linear equations.

### Inspect of data accuracy

The accuracy of nearest-neighbor parameters is our major concern, to inspect the data accuracy, thermodynamic parameters for each of the RNA/DNA hybrid duplexes used in our study were predicted from the determined nearest-neighbor parameters. These predicted parameters were then compared with experimental observed results to inspect the error of prediction and actual value. Linear regression of predicted and experimental values is conducted for predictive analysis. Furthermore, a leave-one-out cross validation (LOOCV) was applied to inspect of data accuracy and avoid overfitting. This is a type of cross-validation approach in which each observation is considered as the validation set and the rest (N-1) observations are considered as the training set. This method helps to reduce bias and randomness. In our LOOCV validation, we conduct the SVD calculation in Equation (15) using all but one thermodynamic parameter from dataset. Nearest-neighbor parameters determined were then used to predict the thermodynamic parameters for the excluded duplex and compare with the experimental data. This process then repeats 227 times to go through the whole dataset. At this step, those duplexes with mismatches less than three-times overdetermined were skipped to reduce the influence of random error from single coverage of data. Finally, the mean squared errors (MSE) were calculated from this process were also compare with those from the full dataset to see how much error would be introduced if a duplex data is unused. This also inspect the ability of accurately predicting thermodynamic parameters of RNA/DNA hybrid duplexes other than our study.

## RESULTS

### Thermodynamic parameters for RNA/DNA hybrid duplexes

The thermodynamic parameters of tested RNA/DNA hybrid duplexes in our study are provided in **Supplementary Table S1**. This table lists both parameters set determined from T_m_^−1^ versus ln C_T_/4 method and Marquardt non-linear least squares curve fit method. For all duplex groups, the average error of ΔG°_37_, ΔH° and ΔS° values from T_m_^−1^ versus ln C_T_/4 method are ± 0.60 kcal/mol, ± 3.95 kcal/mol and ± 11.7 cal/mol·K, from Marquardt non-linear least squares curve fit method are 0.02 kcal/mol, ± 0.25 kcal/mol and 0.80 cal/mol·K, respectively. Linear regression shows a good agreement of ΔG°_37_, ΔH° and ΔS° from two methods with an adjusted R-square of 0.96, 0.95 and 0.96, respectively (**Supplementary Figure S1**). The maximum difference between ΔH° values form two methods is below 15%, validating the two-state approximation. Melting temperatures of all duplexes are in range of 33.3°C to 58.4°C. For the complete dataset with parameters from literature, the average differences between ΔG°_37_, ΔH° and ΔS° values from two methods are 2.99%, 3.93% and 4.39%, respectively. Average coverage for mismatch involving pairs is 3.19 (from 2 to 4), indicating a uniform coverage of mismatch involving pairs. Therefore, we got an exhaustive dataset for approximating nearest-neighbor parameters.

### Comparation of experimental and nearest-neighbor predicted thermodynamic parameters of RNA/DNA hybrid duplexes

A coefficient matrix of 228 rows and 113 columns was constructed from our dataset for SVD calculation. All 113 possible nearest-neighbor parameters determined in our study are listed in **Table 1**. Average estimation errors in ΔG°_37_, ΔH° and ΔS° are ± 0.18 kcal/mol, ± 1.39 kcal/mol and ± 4.32 cal/mol·K, respectively. These nearest-neighbor parameters were used to predict the thermodynamic parameters of duplexes used in our study for a comparation with reported or experimental data. Results of each duplex are listed in **Supplementary Table S4**, it also lists the standard deviations for each of the duplexes containing individual internal mismatches. The average diversion of prediction and experiment data is 0.26 kcal/mol in ΔG°_37_ (3.55%), 5.05 kcal/mol in ΔH° (6.70%), 16.3 cal/mol·K in ΔS° (7.50%) and 1.14°C in T_m_ (2.91%). Linear regression on predicted values versus experimental values on above parameters shows the adjusted R-square of ΔG°_37_, ΔH°, ΔS° and T_m_ to be 0.97, 0.88, 0.87 and 0.97. The slope of each fitted equations are 1.0000 ± 0.0122, 1.0000 ± 0.0244, 1.0000 ± 0.0253 and 0.9714 ± 0.0121, which are all close to the ideal value of 1.0 (Figure 1). These results suggest that the prediction excellently matches the experimental data.

**Table 1.**
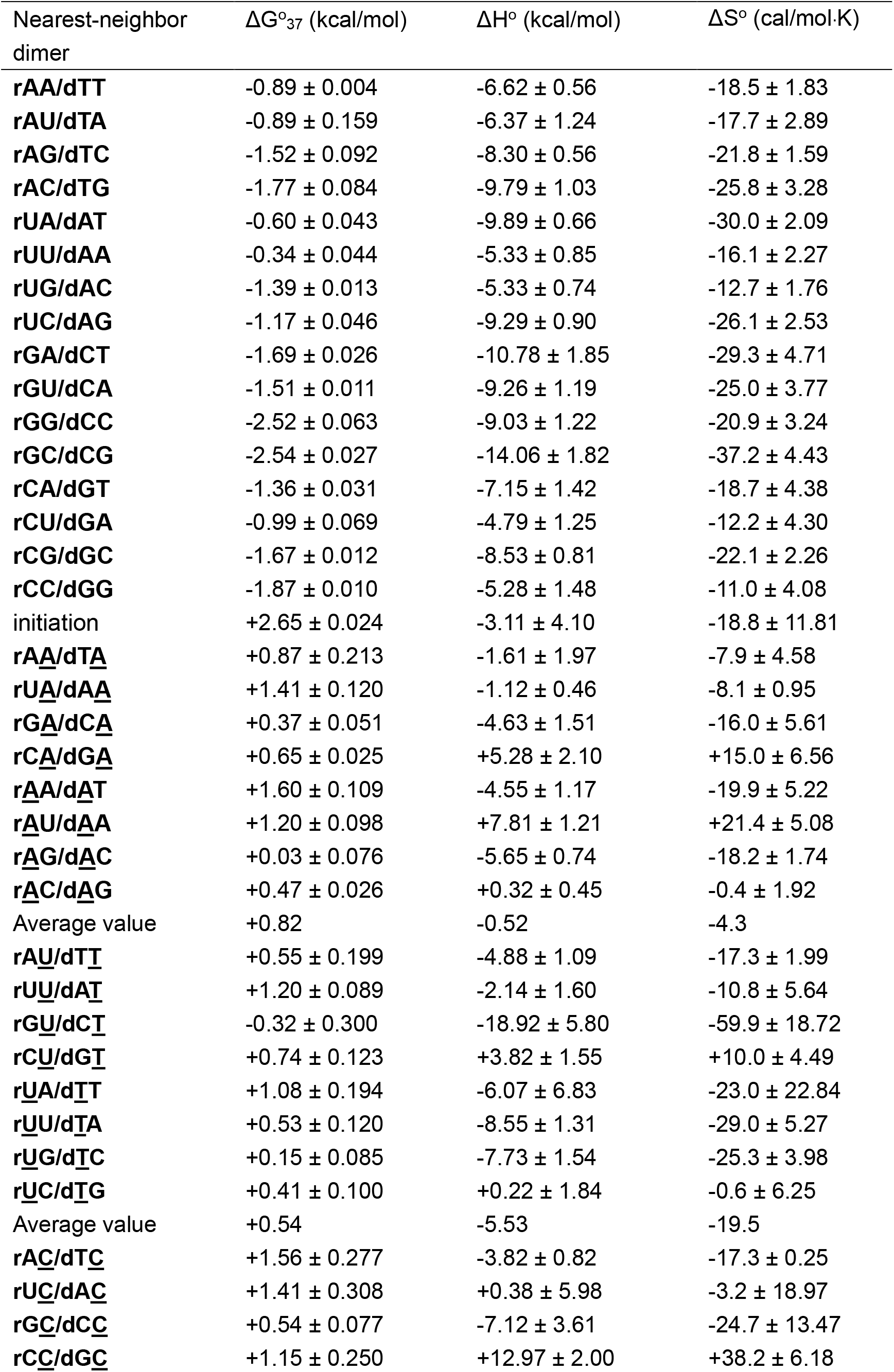
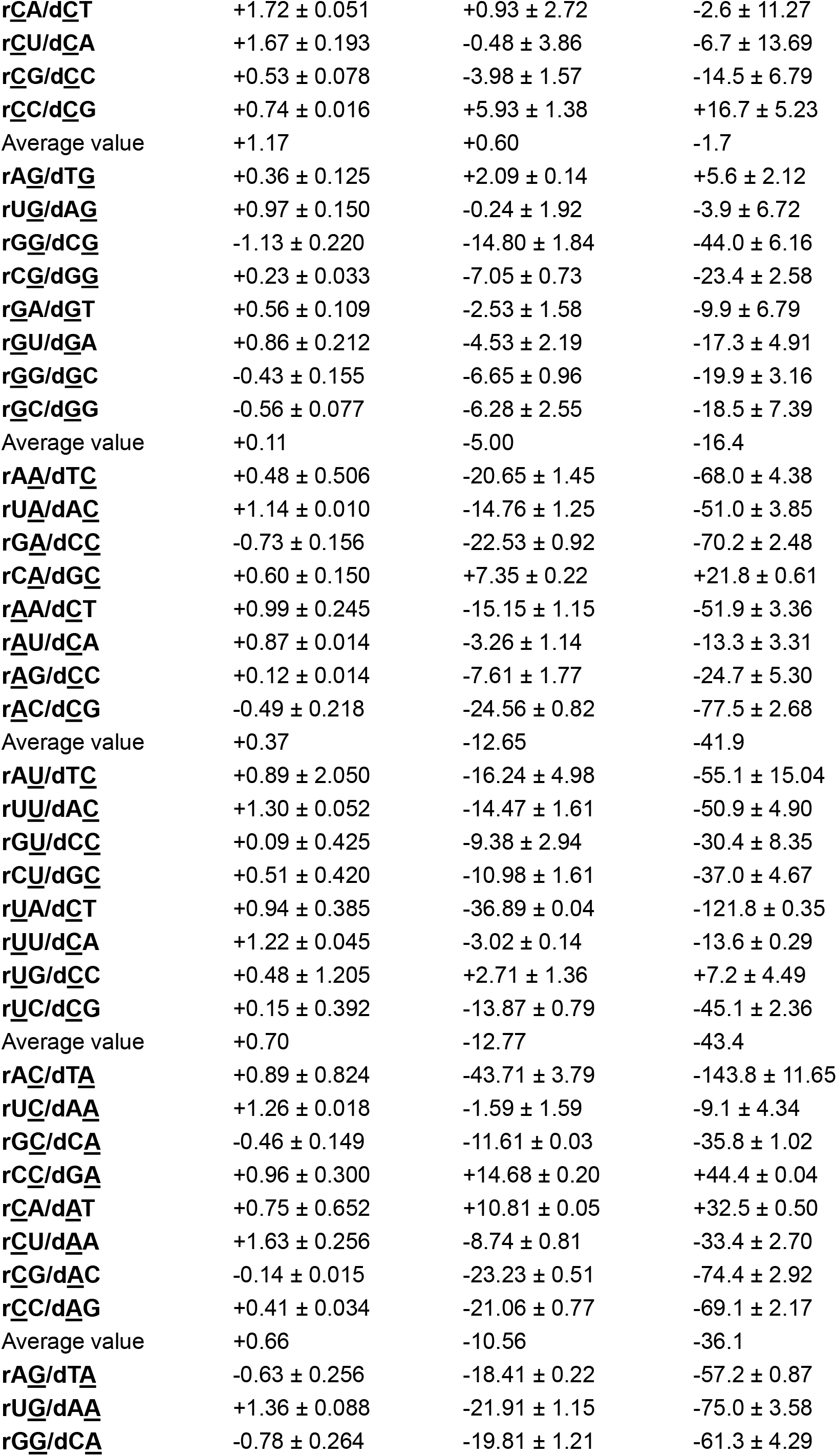
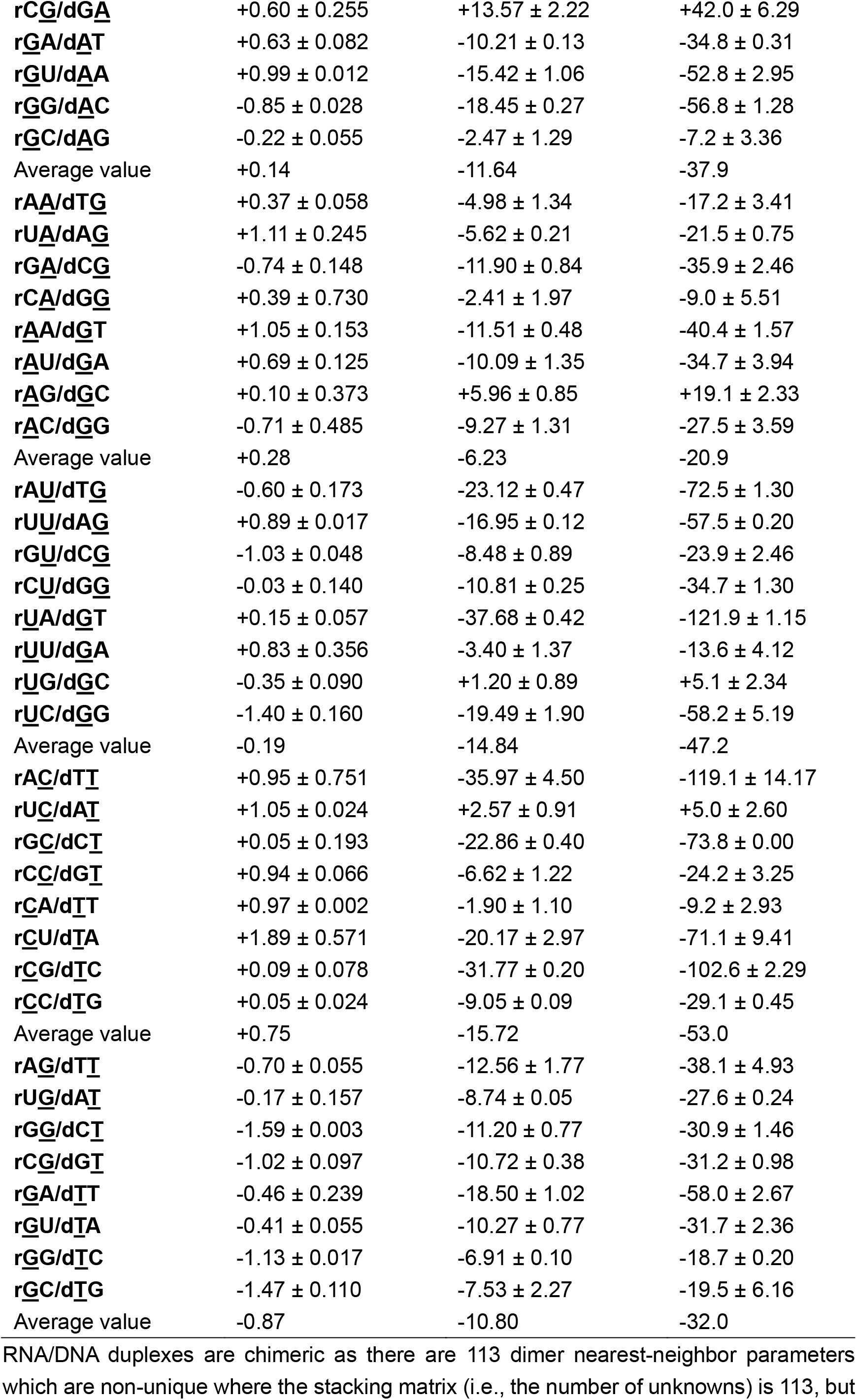
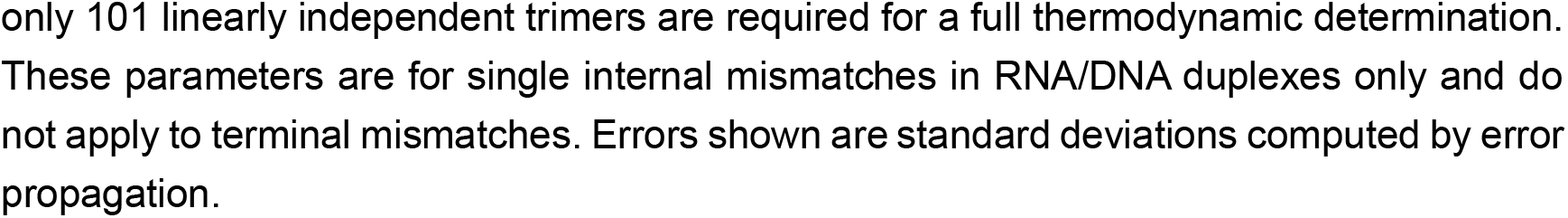
Non-unique nearest-neighbor thermodynamics in RNA/DNA duplexes in 1 M NaCl, 10 mM Na_2_HPO_4_, 0.5 mM Na_2_EDTA, pH 7.0

**Figure 1.**
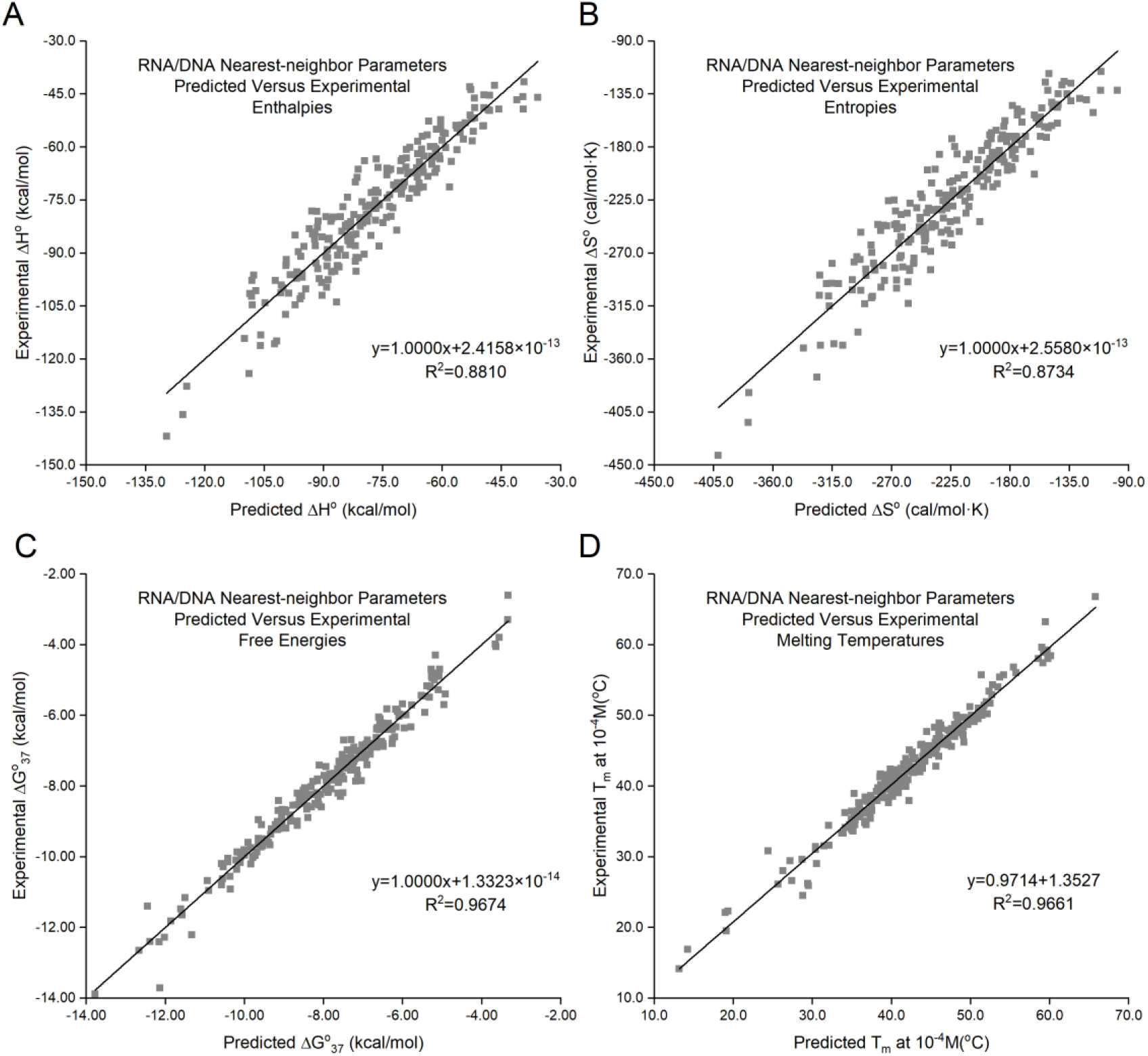
Comparison of experimental versus predicted **(A)** enthalpies; **(B)** entropies; **(C)** free energies and **(D)** melting temperatures for RNA/DNA duplexes with single internal mismatches. Entropies, free energies, enthalpies and melting temperatures are predicted within 5.05 kcal/mol, 16.3 cal/mol·K, 0.26 kcal/mol and 1.14°C, on average, respectively.

### LOOCV validation of data accuracy

The LOOCV validation was conducted for preventing overfitting. It proved the accuracy of determined parameters. The average diversions are 0.45 kcal/mol in ΔG°_37_ (6.17%), 9.13 kcal/mol in ΔH° (11.90%), 29.32 cal/mol·K in ΔS° (13.25%) and 1.97°C in T_m_ (5.03%), respectively. Pearson correlation of these parameters are 0.9542, 0.8213, 0.7969 and 0.9584 (Figure 2, raw data for all duplexes are shown in **Supplementary Table S5**). Further analysis of parameter sets from LOOCV shows good agreement with parameters in Table 1 with an average variance of ΔG°_37_, ΔH° and ΔS° are 0.02, 0.42 and 1.36, respectively (**Supplementary Table S6**). This indicates that lacking any duplex in our study would have no strong impact on the final determined nearest-neighbor parameters, proving the robustness of our study. As a result, nearest-neighbor parameters from our study have excellent accuracy in predicting the thermodynamic parameters of RNA/DNA hybrid duplexes unseen.

**Figure 2.**
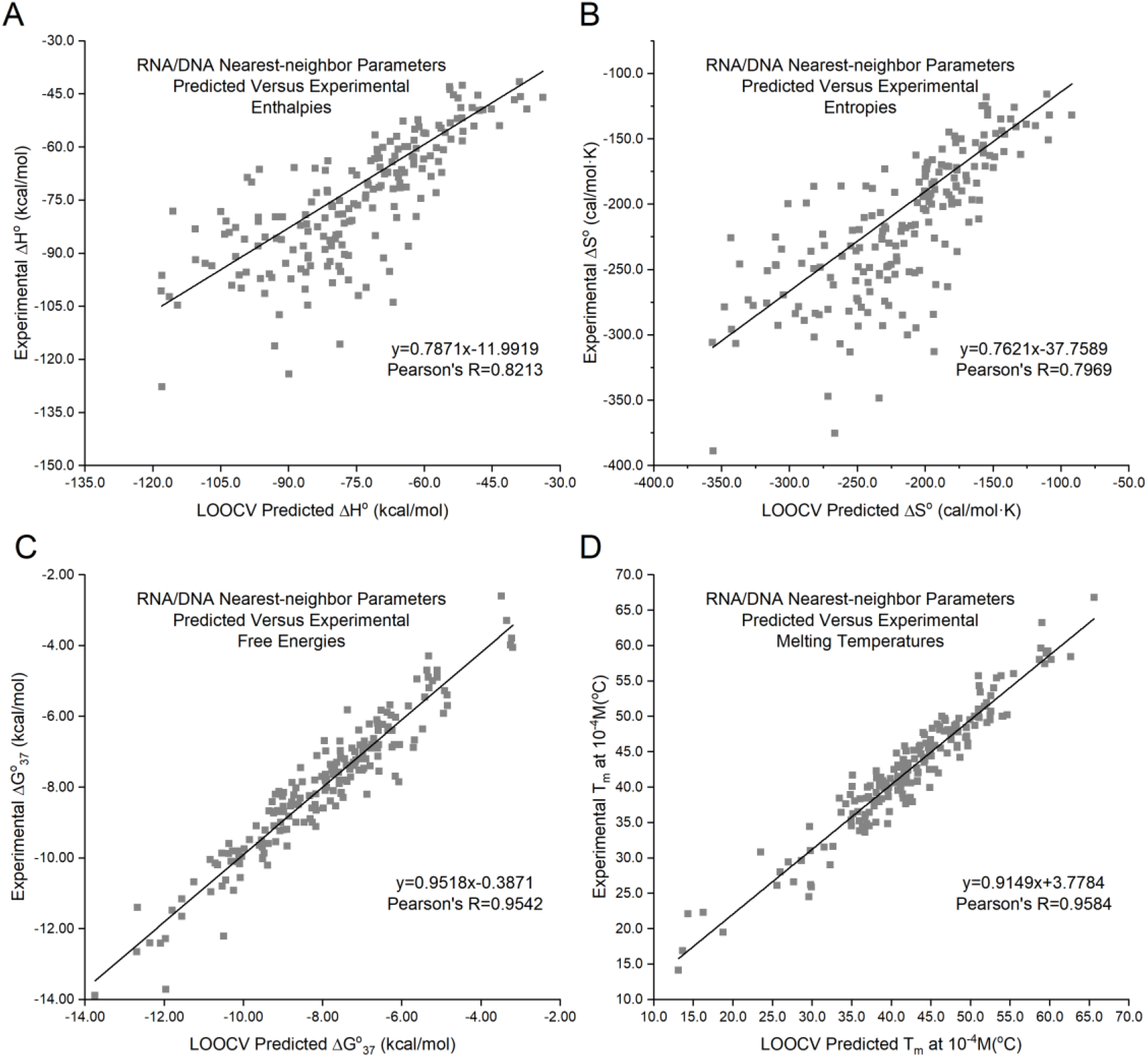
Comparison of experimental versus LOOCV predicted **(A)** entropies; **(B)** enthalpies; **(C)** free energies and **(D)** melting temperatures for RNA/DNA duplexes with single internal mismatches. Entropies, free energies, enthalpies and melting temperatures are predicted within 9.13 kcal/mol, 29.32 cal/mol·K, 0.45 kcal/mol and 1.97°C, on average, respectively.

### Trends of stability of Watson-Crick nearest-neighbor pairs

All ΔG°_37_ for nearest-neighbor parameters of Watson-Crick base pairs are below zero, proving the stability of Watson-Crick base pairs under 37°C. The order of observed stability is rGC/dCG ≈ rGG/dCC > rCC/dGG > rAC/dTG > rGA/dCT ≈ rCG/dGC > rAG/dTC ≈ rGU/dCA > rUG/dAC ≈ rCA/dGT > rUC/dAG > rCU/dGA > rAU/dTA ≈ rAA/dTT > rUA/dAT > rUU/dAA, this trend suggests that both base composition and sequence are important determinants of RNA/DNA duplex stability. The most stable Watson-Crick base pair is rGC/dCG, and stability drops with the composition of A/T and U/A pairs. This consistent with the property that G/C base pairs are more stable than A/T base pairs for differences in hydrogen bonding and stacking. Moreover, we found the stability of rUU/dAA is significantly smaller than other Watson-Crick base pairs, which is consist with previous report that rU/dA sequences are exceptionally unstable (72). On the other hand, the nearest-neighbor ΔH° and ΔS° parameters have no similar trend.

### Trends of stability of nearest-neighbor pairs involving mismatches

For nearest-neighbor pairs involving mismatches, literatures suggest that the stabilities were significantly affected by both the identity of the mismatch and the adjacent Watson-Crick base pairs (54–60,73). The nearest-neighbor free energy and enthalpy contributions for single nucleotide internal mismatches of both 5’- and 3’-contexts have been compared using Table 1 and Equation (16)

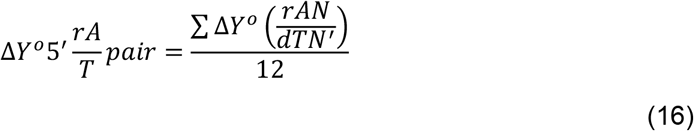

Where Y = G, H or S.

The stability trend for the base pairs 5’-of the single internal mismatch is rG/dC >> rA/dT ≈ rC/dG > rU/dA with the average ΔG°_37_ contribution of −0.48, +0.42, +0.48, +1.08 kcal/mol, respectively. The 3’-of the single internal mismatch is rC/dG > rG/dC >> rA/dT > rU/dA with the average ΔG°_37_ contribution of −0.21, −0.11, +0.83, +1.00 kcal/mol, respectively, which is a consistent result of G/C pairs has higher stability than A/T pairs for more hydrogen bonds and differences in stacking.

Our result shows that differences of the stability of nearest-neighbor pairs involving mismatches at 37 °C (ΔG°_37_) were ~2 kcal/mol depending on the mismatch base pairs. The order of average stabilities for mismatches is rG/dT > rU/dG > rG/dG > rG/dA > rA/dG > rA/dC > rU/dT > rC/dA ≈ rU/dC ≈ rC/dT > rA/dA > rC/dC, with average ΔG°_37_ of −0.87, −0.19, +0.11, +0.14, +0.28, +0.37, +0.54, +0.66, +0.70, +0.75, +0.82 and +1.16 kcal/mol, respectively. For mix-bases the trend is rG/dD > rU/dB > rA/dV > rC/dH (average ΔG°_37_ are −0.21, +0.35, +0.49, +0.86 kcal/mol). The results also prove that for RNA/DNA hybrid duplexes, pairs with the same mismatched nucleobases pairs but positions are exchanged should be considered to be different in thermodynamics (for example, G/A mismatches can be constructed for rG/dA and rA/dG pairs). All ΔG°_37_ for nearest-neighbor parameters involving mismatches are above zero except two: rG/dT and rU/dG. This result shows that rG/dT and rU/dG mismatches both enhance the duplex stability. rG/dT has the highest stability among all mismatch involving pairs, even have lower ΔG°_37_ than Watson-Crick pair rUU/dAA and rUA/dAT. Result shows that rG/dT containing duplexes were always more stable than those of rU/dG, which is consistent with literature reports for single internal mismatches in RNA and DNA duplexes (73). These data support the hypothesis that a nearest-neighbor context dependence exists for single internal mismatches in RNA/DNA hybrid duplexes (60,73).

### Trends of stability of trimers involving mismatches

According to nearest-neighbor model, we can calculate the contribution of a trimer to an RNA/DNA duplex by adding up the parameters of its nearest-neighbor dimers. Therefore, a full set of ΔG°_37_ parameters of the 12 mismatch involving trimers with 5’- and 3’-Watson-Crick closing pairs is calculated and listed in **table 2**. Among all trimers, the most stable trimer is rGGC/dCTG with the ΔG°_37_ of −3.06 kcal/mol while the most unstable trimer is rACA/dTCT, which the ΔG°_37_ is +3.28 kcal/mol. The rG/dT is also the most stable mismatch, the ΔG°_37_ values ranging from −0.58 (rUGU/dATA) to −3.06 kcal/mol (rGGC/dCTG). All of the 16 trimers involving rG/dT have the lowest ΔG°_37_ with the same 5’- and 3’-Watson-Crick closing pairs ranging from −3.06 (rGGC/dCTG) to −0.58 (rUGU/dATA) kcal/mol. rU/dG involving trimers are the second most stable mismatch type with the ΔG°_37_ range from 1.72 (rUUU/dAGA) to −2.44 kcal/mol (rGUC/dCGG), 12 out of total 16 trimers involving rU/dG have the second lowest ΔG°_37_ with the same 5’- and 3’-Watson-Crick closing pairs. rC/dC is destabilizing in all contexts with ΔG°_37_ values ranging from 1.07 (rGCG/dCCC) to 3.28 kcal/mol(rACA/dTCT). For all contexts rC/dC is less stable than other mismatches except for rCXU/dGXA, in which rCCU/dGTA is only 0.01 kcal/mol higher in ΔG°_37_. The relative stability of other mismatch involving trimers is appear to be context dependent (Figure 3), and a general trend of rG/dT > rU/dG > rG/dG ≈ rG/dA > rA/dG > rA/dC > rU/dT > rC/dA ≈ rU/dC ≈ rC/dT > rA/dA > rC/dC. This matches the trend in nearest-neighbor parameters.

**Table 2.**
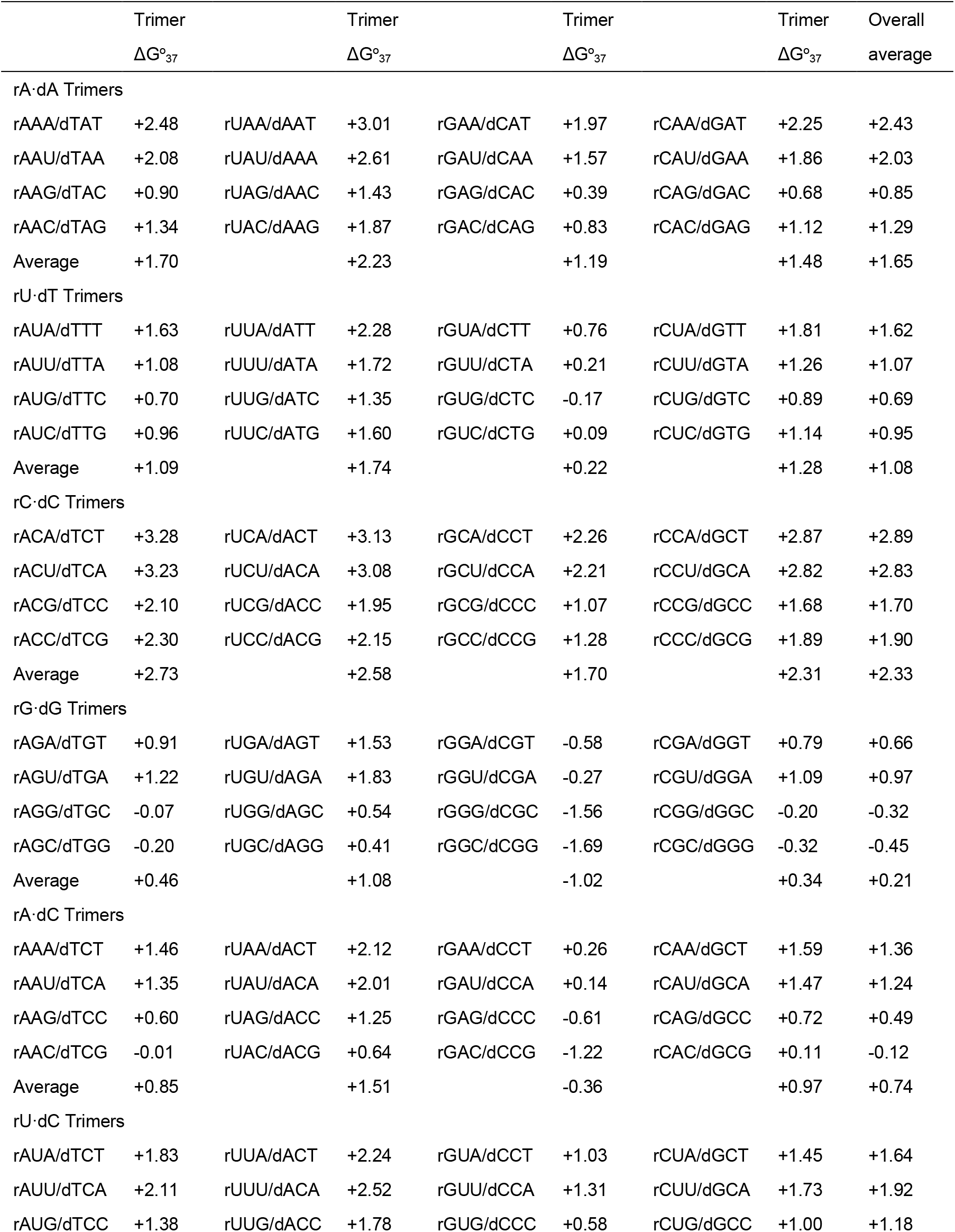
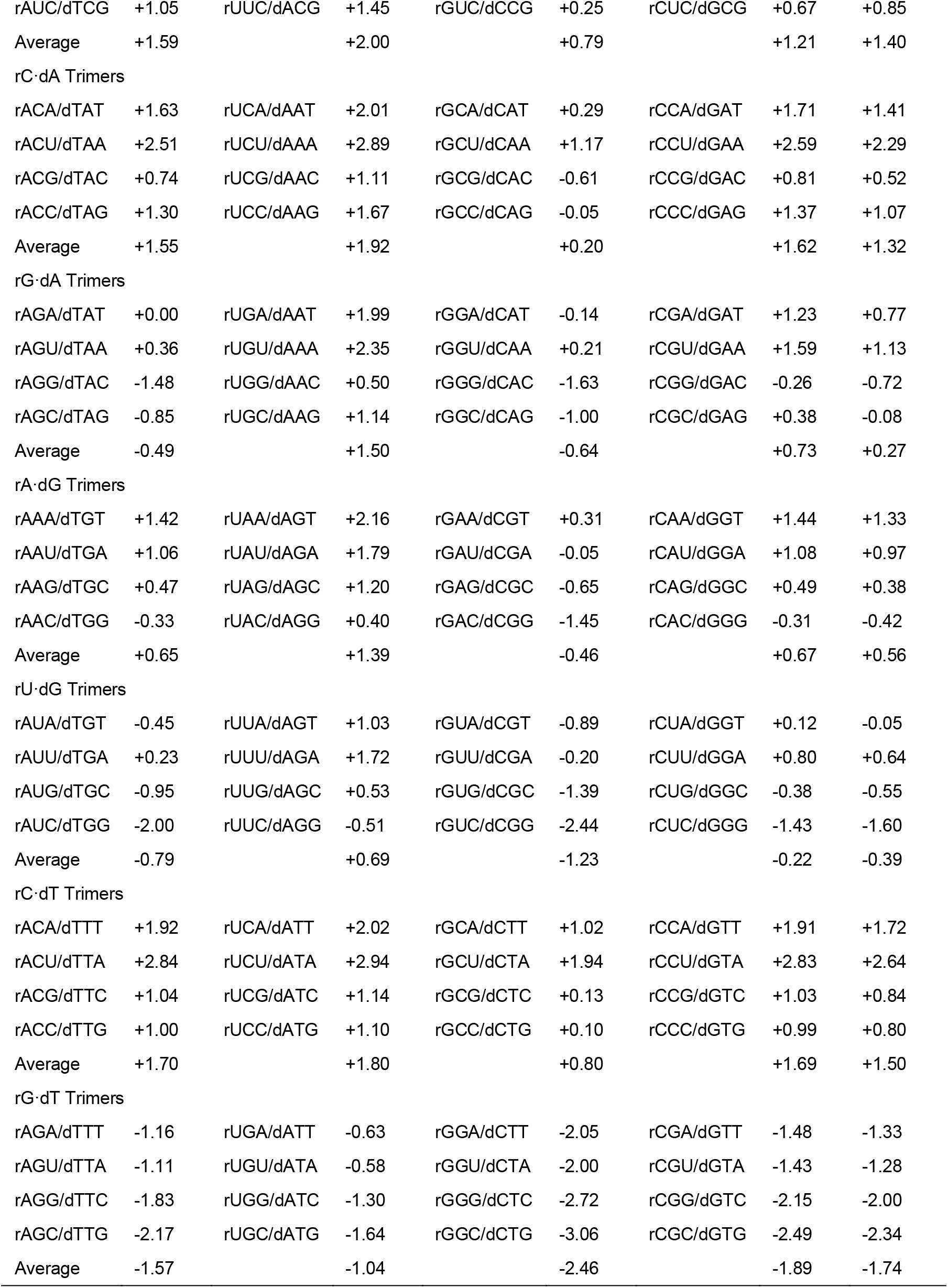
The 16 trimer combinations for each of the reported single internal mismatches in RNA/DNA duplexes. These values were calculated using the parameters in Table 1

**Figure 3.**
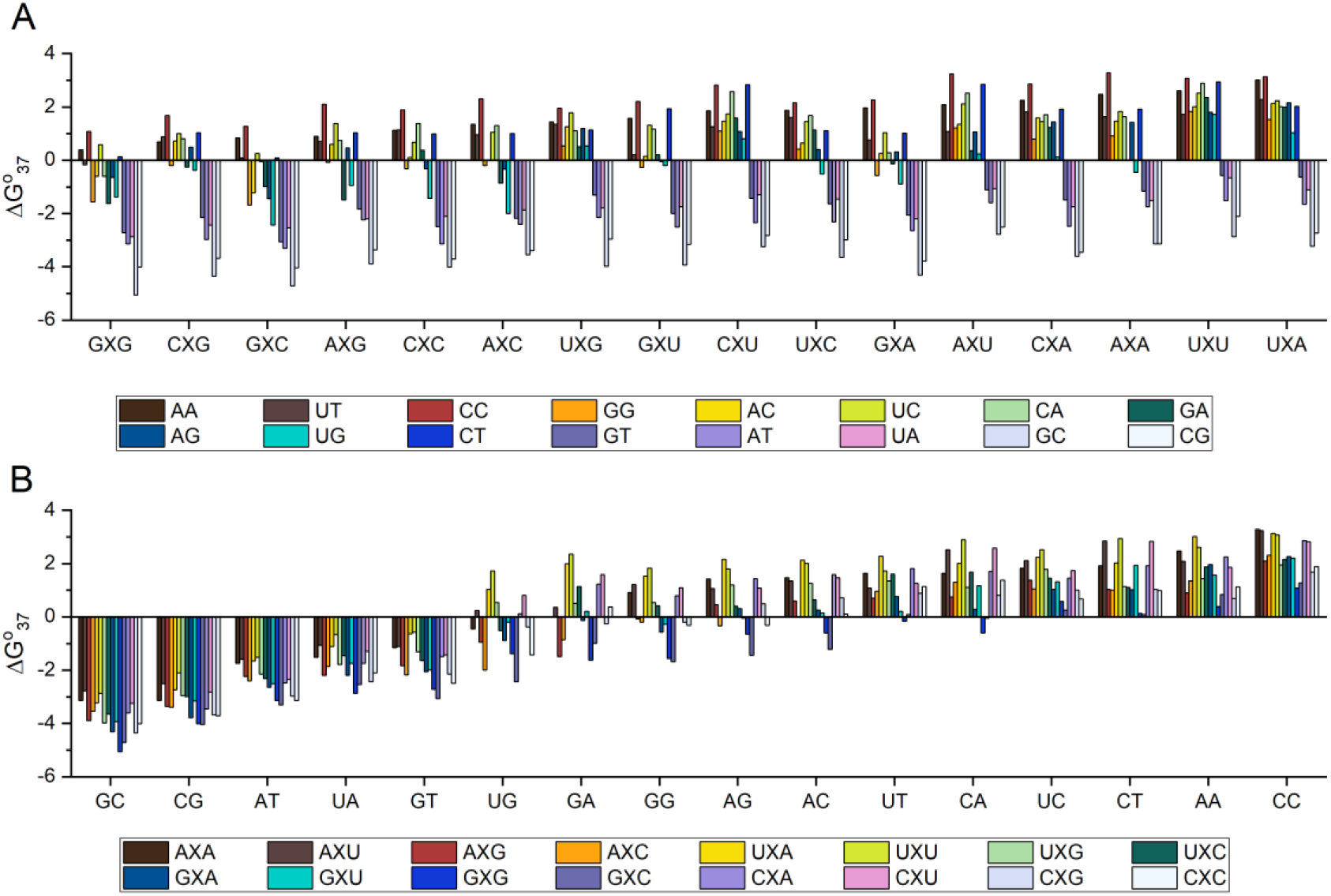
Trends of stability of trimers involving mismatches. Trends of different **(A)** mismatches and **(B)** 5’- and 3’-adjacent Watson-Crick base pairs. The general trend was rG/dT > rU/dG > rG/dG ≈ rG/dA > rA/dG > rA/dC > rU/dT > rC/dA ≈ rU/dC ≈ rC/dT > rA/dA > rC/dC.

### Structural studies on stability of nearest-neighbor pairs involving mismatches

Rational explanation for our observation on mismatch stability could be found in oligonucleotide structural studies. We conclude that the stability trend has a combined pattern, which is determined by number of hydrogen-bonds between different base pairs and influenced by stacking interactions. A decrease of average predicted H-bond numbers can be observed follow the stability trend. For the most stable rG/dT and rU/dG mismatches (**Figure 4**), literature shows both mismatch types form similar wobble hydrogen-bonded structure in diverse contexts (54), and a possible structure of three H-bonds with enol tautomer of T and U (74,75). This could contribute to the extra stability of these two mismatches. Additional stability of rG/dT is most likely due to the different effects of the 2’-hydroxyl group in ribose for the lacking of C5 methyl in uracil (73). For the rest of mismatches, rG/dG, rG/dA, rA/dC, rA/dG, rU/dT, rC/dA, rU/dC and rC/dT are all reported to have possible structure with two H-bonds (55–58) (**Figure 4**), and the most unstable rA/dA and rC/dC pairs have only one H-bone structure.

**Figure 4.**
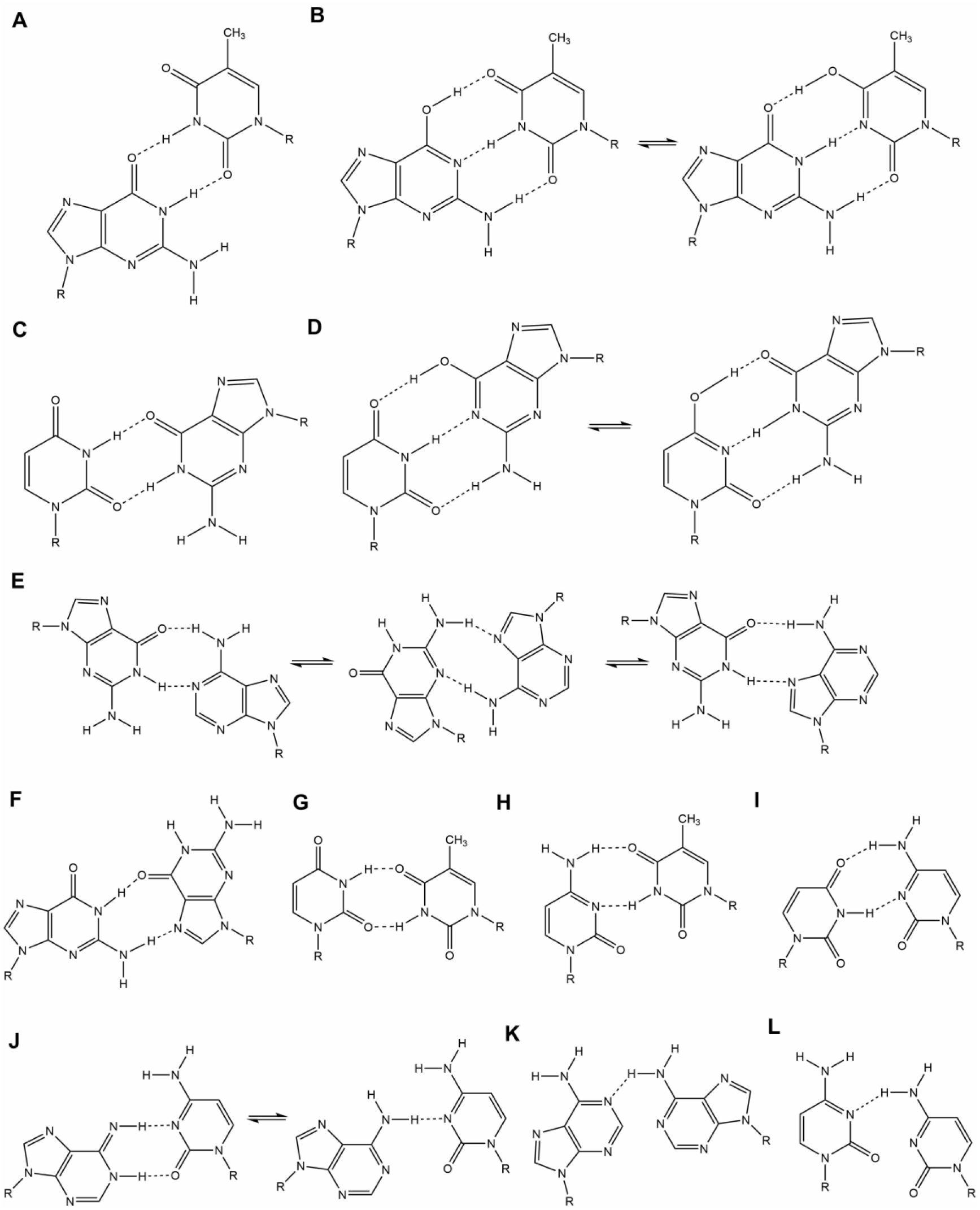
Proposed structures for single internal mismatches in RNA/DNA duplexes. The structures are **(A)(B)** rU/dG; **(C)(D)** rG/dT; **(E)** rG/dA and rA/dG; **(F)** rG/dG; **(G)** rU/dT; **(H)** rC/dT; **(I)** rU/dC; **(J)** rA/dC and rC/dA; **(K)** rA/dA **(L)** rC/dC.

Another possible contribution to the stability trend is the base stacking. rG/dG, rG/dA and rA/dG mismatches, which are the most stable three types among all mismatches with two H-bonds, have a high stacking potential due to their purine rings while the conformation does not perturb the helical backbone structure of the duplex (57). This also works for rA/dA but the short of H-bond reduces the stability. rU/dT and rC/dC mismatches on the other hand, have a relatively lower stacking potential for their pyrimidine ring. These two factors can explain the trend of stability of mismatches we observed: stability is major effected by hydrogen-bonding and minor effected by base stacking in RNA/DNA hybrid duplexes.

### Comparison of Watson-Crick nearest-neighbor parameters of RNA/DNA hybrid duplexes with those in RNA/RNA and DNA/DNA duplexes

SantaLucia *et al*. and Freier *et al*. reporeted the nearest-neighbor parameters of DNA/DNA and RNA/RNA duplexes (33,76). These values are come from the measured thermodynamic parameters and is independent of the duplex sequences. In both DNA/DNA and RNA/RNA parameters, the initiation ΔH° are both assumed to be zero (33,34) while our result shows that in RNA/DNA hybrid duplexes is −3.11 kcal/mol. The ΔS° of initiation is −18.84 cal/mol·K, lower than both in DNA/DNA (−5.90 cal/mol·K for dG/dC and −9.00 cal/mol·K for dA/dT) and RNA/RNA (−10.80 cal/mol·K) duplexes. The fitted ΔG°_37_ of initiation in RNA/DNA is +2.64 kcal/mol, close to those in DNA/DNA (+1.82 kcal/mol for dG/dC and +2.8 kcal/mol for dA/dT) and lower than RNA/RNA (+3.40 kcal/mol). These nearest-neighbor parameters show that the enthalpy contribute to the formation of hybrid duplex, while the entropy is a drawback of duplex formation. This is probably because of the energy change during the formation of H-bonds. The ΔG°_37_ of initiation is above zero and significantly higher than other nearest-neighbor dimers, showing that the formation of first base pair in RNA/DNA hybrid duplexes is much more difficult than extension of duplexes. This proves that the formation of the first base pairs is the most important step in RNA/DNA hybrid duplexes formation.

For other nearest-neighbor dimers, the sequence stability trend in RNA/DNA duplex has a similar pattern as in DNA/DNA and RNA/RNA duplexes, sequences with higher GC content have higher stability. The average stability (ΔG°_37_) of RNA/DNA and DNA/DNA dimers are close (−1.42 and −1.38 kcal/mol), both are strongly lower than RNA/RNA dimers (−1.92 kcal/mol). The RNA/RNA duplex is the most stable among the three kinds of dimers with the same nearest-neighbor sequences. For dimers with same nearest-neighbor sequences in RNA/DNA and DNA/DNA duplex, which one is more stable has no clear patten and is depended on the sequence. For example, rGA/dCT is more stable than dGA/dCT (ΔG°_37_ are −1.68 and −1.46 kcal/mol), but dCG/dGC is more stable than rCG/dGC (ΔG°_37_ are −2.09 and −1.67 kcal/mol). The average ΔH° and ΔS° of RNA/DNA duplex are −8.11 kcal/mol and −21.57 cal/mol/K, both are larger than the average ΔH° and ΔS° of RNA/RNA duplex (−9.80 kcal/mol and −25.38 cal/mol/K) and lower than DNA/DNA duplex (−7.74 kcal/mol and −20.44 cal/mol/K). The trend of ΔH° and ΔS° in these three types of duplexes is depended on the exact sequence and has no clear patten.

### Comparison of single internal mismatches in RNA/DNA hybrid duplexes with DNA/DNA duplexes

The whole set of nearest-neighbor parameters with single internal mismatches in RNA/DNA duplex are compared with those parameters with the same nearest-neighbor sequences in DNA/DNA duplexes reported in literature (54–58). The average ΔG°_37_ of mismatch pairs is +0.37 kcal/mol in RNA/DNA duplex, lower than in DNA/DNA (+0.42 kcal/mol). Peyret *et al*. (57) reported a stability trend of dG/dG > dT/dT > dA/dA > dC/dC (average ΔG°_37_ are −2.23, +0.42, +0.48 and 0.97 kcal/mol, respectively). This trend is identical in RNA/DNA as the average ΔG°_37_ of internal rG/dG, rU/dT, rA/dA and rC/dC mismatches are +0.11, +0.54, +0.82 and +1.16 kcal/mol. The exact other of stability has no clear trend and is depended on the sequence. When comparing stability of mismatches in RNA/DNA and DNA/DNA duplexes, which is more stable is depended on the exact sequence and highly correlated to base pair structure. Mismatches involving two purine rings (G/G, G/A and A/A) are more stable in DNA/DNA duplex. At the same time, mismatches involving one purine (G/T, G/U and A/C) are more stable in RNA/DNA duplex. For the mismatches involving no purine (U/T, T/T, C/T, C/U and C/C), the stability in RNA/DNA and DNA/DNA are close and no clear trend is observed. We believe this can be rationalized by considering the differences in stacking due to the variation of helical structure of nucleic acid chains. In DNA duplexes, the backbone exhibit a B-form conformation characterized by a C2’-endo sugar conformation (77), a weak stacking is observed in this structure (33,78). However, when RNA and DNA are annealed together to form a hybrid duplex, the RNA usually remain A-form C3’-endo conformation, and the DNA usually assume an equilibrium between the C2’ and the C3’-endo conformations (79,80). This conformational equilibrium may interfere with the contribution of the base stacking energies to the hybrid duplex (60), So that mismatches with stronger stacking interactions, like G/G, G/A and A/A, would have higher stability in DNA/DNA duplexes. As for mismatches involving no purine, this type of mismatch has the smallest molecular size that they are not cooperate well with both A-from and B-from backbone structure. This could explain why their stability are close in these two types of duplexes.

When considering internal mismatched trimer stability, the most and least stable trimer in DNA/DNA duplex is dGGC/dCGG and dACT/dTCA, which contribute to total ΔG°_37_ from −2.22 to +2.66 kcal/mol, respectively. RNA/DNA duplex has a wider range of trimer contribution from −3.06 to +3.28 kcal/mol, the upper and lower limits are rACA/dTCT and rGGC/dCTG, both are different to DNA/DNA duplex. Trend of the effect on stability of 5’-adjacent Watson-Crick base pairs are similar in RNA/DNA and DNA/DNA duplexes: G/C are most stable and U(T)/A are least stable, but effect of 3’-adjacent pairs are significantly different. In DNA/DNA duplexes, this trend is C > G > U > A. Unlike in RNA/DNA duplex which the 3’-G and C trimers has much greater stability than U and A, such great gap is not observed in DNA/DNA duplex for 3’-G and C trimers comparing to T and A.

### Comparison of single internal mismatches in RNA/DNA hybrid duplexes with RNA/RNA duplexes

Freier *et al*. (76) reported nearest-neighbor parameters of single internal mismatches in RNA/RNA duplex. The average ΔG°_37_ of mismatch pairs is +0.37 kcal/mol in RNA/DNA duplex, significantly higher than in RNA/RNA (−1.10 kcal/mol). Each mismatch is more stable in RNA/RNA duplexes than in RNA/DNA hybrid duplexes for ~1.47 kcal/mol. When comparing the stability trend of mismatches from two types of duplexes, we discover the mismatches can be divided into two parts: (i) mismatches involving two purine rings and (ii) other mismatches. When comparing two parts separately, the stability trends of mismatches in these two parts are identical in RNA/DNA and RNA/RNA duplexes, while mismatches involving two purine rings have higher stability rank in RNA/RNA duplex than in RNA/DNA duplex. This can be similarly explained by stacking. In RNA/RNA duplex, the nucleic acid backbone exhibit an A-form conformation characterized by a C3’-endo sugar conformation (81). In this kind of conformation base pairs arranges tight to each other therefore strengthen the stacking contribution (82). We believe this conformation added the stability of mismatches with two purine rings and increased the stability of G/G and G/A mismatches.

For trimer stability, the most and least stable trimer in RNA/RNA duplex are rCGC/rGUG and rUUA/rAUU with the ΔG°_37_ from −3.7 to −1.0 kcal/mol, both are different from RNA/DNA. Trimers of RNA/RNA are more stable than in RNA/DNA duplex with an average gap of 2.95 kcal/mol, while the difference in free energies in RNA/RNA compared to RNA/DNA hybrid duplexes is sequence specific: rU/dG and rU/rG has the second closest stability of 2.34 kcal/mol in difference while rA/dA and rA/rA has the largest difference of 4.05 kcal/mol. The difference of ΔG°_37_ between rG/dT and rG/rU is 0.99 kcal/mol, this value is the lowest among all mismatches and significantly lower than average gap. This proves again the rG/dT mismatch has additional stability. Again, the rationalization for this observation was based upon the C5 methyl affection on 2’-hydroxyl ribose in thymine.

### Comparison with other parameters from literature

Nearest-neighbor parameters for Watson-Crick pairs and part of internal mismatch pairs in RNA/DNA hybrid duplexes have been determined in literature (35,60). All parameters from these two researches are determined by experimental data which also incorporated into dataset of our study. For comparation, we not only compared parameters from our study with previous works, but also conducted the SVD calculation in dataset of these works to further validate the parameters and calculation method.

Sugimoto *et al*. reported nearest-neighbor parameters of Watson-Crick pairs. There is a large difference between our parameters and theirs. For example, the rCA/dGT has ΔG°_37_ of −1.36 kcal/mol in our parameters, but −0.9 kcal/mol from Sugimoto’s work. The average differences of our and Sugimoto’s parameters on ΔG°_37_, ΔH° and ΔS° are 18.9%, 51.6% and 67.8%, respectively. However, our parameters have better prediction on experimental thermodynamic data. The average differences between predicted and experimental values in Sugimoto’s work are 5.09% for ΔG°_37_, 7.01% for ΔH°, 7.95% for ΔS° and 4.64% for T_m_, all are larger than our result (3.55% in ΔG°_37_, 6.70% in ΔH°, 7.50% in ΔS° and 2.91% in T_m_). We further conducted SVD calculation on duplexes used in Sugimoto’s work, the resulted parameters predict experimental thermodynamic data with errors of 3.83% in ΔG°_37_, 5.95% in ΔH°, 6.70% in ΔS° and 3.74% in T_m_, respectively. This result indicates that our algorithm has better performance comparing with the algorithm used in Sugimoto’s work.

Watkins *et al*. reported 32 nearest-neighbor parameters involving rA/dA, rC/dC, rG/dG and rU/dT four types of mismatches (60). Strong differences also found in our and their parameters. Since some parameters have low absolute value (for example, rUC/dTG from Watkins’ work is −0.02), the difference ratio cannot represent well with the difference. So we use the prediction error to compare the two sets of parameters. In Watkins’ work, the parameters are determined from a different way by subtracting the core sequence from experimental sequence to get the trimer parameters. It is based on Sugimoto’s work and used their parameters to calculate trimer parameters. So, we use a combination of Watkins’ parameters and Sugimoto’s parameters to predict sequences thermodynamic parameters. The resulted errors are 6.05% in ΔG°_37_, 10.00% in ΔH°, 11.04% in ΔS° and 5.50% in T_m_, about 1.5-fold higher than our prediction error. Thus, our parameters should give improved predictions for the stability of RNA/DNA hybrid duplexes.

## DISCUSSION

In this work, we measured the melting curves of 99 RNA/DNA hybrid duplexes with internal mismatches and determined their thermodynamic parameters by two separate ways. A full set of 113 nearest-neighbor parameters were determined based on nearest-neighbor model using singular value decomposition from a combination of our results with literature reported thermodynamic parameters (35,60). Determined nearest-neighbor parameters accurately predicted thermodynamic parameters of the RNA/DNA duplexes. We first introduced data validation progress in similar research, the leave-one-out cross validation, and the result proved the robustness of our study. Our parameters have the best reported prediction accuracy when comparing with other parameters from literature. Hence, we developed a full set of nearest-neighbor parameters that can accurately predict the thermodynamic parameters of RNA/DNA hybrid duplexes. Furthermore, we analyzed the stability trend of nearest-neighbor pairs and found a mismatch pair related pattern, we realized that both hydrogen bonding and base stacking contribute to the stability of mismatches, the stability is mainly depended on the number of hydrogen-bonds and secondary depends on the stacking between pairs. Moreover, we compared the nearest-neighbor sequences in RNA/DNA hybrid duplexes with DNA/DNA and RNA/RNA duplexes, and found a structure depended pattern of stability. In conclusion, this work can help to thermodynamically predict the properties of RNA/DNA hybrid duplexes.

Ordinary researches on nearest-neighbor models determine the nearest-neighbor parameters from experiments, and validate the parameters using the same sequences. In other words, the training and testing sets are the same. Since parameters naturally perform well on testing sets already seen, this method has potential inability in predicting duplexes unseen. Inspired by cross validation in machine learning, we introduced data validation process in our study: a LOOCV validation, which is the first time to introduce data validation in similar researches. Under data validation, the correlation between predicted and experimental ΔG°_37_ and T_m_ are significantly higher than ΔH° and ΔS°. One possibility of cause is the error. ΔG°_37_ and T_m_ have lower error than ΔH° and ΔS°, thus even under same coverage, the accuracy of LOOCV validation could be different. One potential point to further improve this validation is the coverage of mismatch pairs, typical coverage of designed sequences is above 2 for mismatch pairs (57,60,76), this coverage could be improved. Besides, melting curve from different equipment could have minor differences even for the same duplexes, since our dataset combines thermodynamic parameters from the literature, another potential improvement could be re-exam of the reported duplexes.

The nearest-neighbor parameters involving mismatches are diverse in free energy and thus result in different stability. During binding, the binding energy differences would result in different binding activities. This indicates that our parameters have potential usage in applications involving RNA/DNA combination, like CRISPR/Cas systems which depended upon the recognize and cleave of RNA and DNA (83). Designed CRISPR guide RNA (crRNA) turns program Cas nucleases to target DNA sequences in which nucleic acid sequence is complementary with the guide portion of the crRNA and proximal to a PAM (protospacer adjacent motif) site. However, binding to unexpected loci sometimes occur, leading to unpredictable results. This phenomenon, referred to as “off-target”, to a big extent, is due to the fact that the binding process can bear several mismatches or bulges between sgRNA and target DNA (84,85). In the design of crRNA, avoiding off-target effects is regarded as the major concern for experimental accuracy. Yet the in-ability to properly account for the effects of nucleotide mismatches often times results in a reduction of target specificity due to the off-target effect. In terms of crRNA designing, while multiple empirical models or principles have been developed based on mismatch number and location (86,87), there are no such models account for mismatch types. Considering the stability differences between different mismatches in this work, like the special stability of rG/dT mismatch, such kind of design principle is inability in avoiding off-targets. Our parameters provide a potential thermodynamic approach of predicting the binding activities of crRNA and off-target sites. We can use thermodynamic parameters from nearest-neighbor model as a direct indicator for off-target efficiency in future works.

## ACKNOWLEDGEMENT

We also thank members from the C. Zhang and H. Xing lab oratories for critical discussions of this work.

