## Supplementary Table S1 for "Thermodynamic Parameters Contributions of Single Internal Mismatches In RNA/DNA Hybrid Duplexes"

| **Table S1:** Thermodynamics of duplex formation of RNA/DNA Hybrid Duplexes containing internal mismatches. All Tm’s are calculated using a duplex concentration of 1e^-4^M. | | | | | | | |
| --- | --- | --- | --- | --- | --- | --- | --- |
| **RNA/DNA Duplex** | **1/Tm Method Curve Fit Method** | | | | | | |
| **A/C Internal Mismatch** | **ΔG°_37_ (kcal/mol)** | **ΔH° (kcal/mol)** | **ΔS° (cal/K*mol)** | **T_m_ (°C)** | **ΔG°_37_ (kcal/mol)** | **ΔH° (kcal/mol)** | **ΔS° (cal/K*mol)** |
| GUUGAACCUAC  CAACTCGGATG | -7.71 ± 0.76 | -124.17 ± 5.92 | -375.50 ± 17.94 | 40.0 | -7.75 ± 0.001 | -124.19 ± 0.002 | -375.46 ± 0.01 |
| GUGUAAGUACC  CACATCCATGG | -7.11 ± 0.55 | -91.90 ± 4.22 | -273.39 ± 12.60 | 39.0 | -7.49 ± 0.01 | -91.55 ± 0.08 | -271.03 ± 0.26 |
| GAUCAAUCCAC  CTAGTCAGGTG | -7.52 ± 0.97 | -84.17 ± 5.54 | -247.13 ± 16.37 | 40.7 | -7.62 ± 0.02 | -83.48 ± 0.10 | -244.59 ± 0.31 |
| CGACCAACUUG  GCTGGCTGAAC | -7.47 ± 0.48 | -66.71 ± 3.18 | -191.00 ± 9.20 | 41.4 | -7.55 ± 0.14 | -64.93 ± 1.19 | -184.98 ± 3.78 |
| UCACACGAUAA  AGTGCGCTATT | -6.69 ± 0.88 | -86.16 ± 5.89 | -256.22 ± 17.62 | 37.6 | -7.01 ± 0.02 | -83.44 ± 0.16 | -246.45 ± 0.52 |
| GCGUCAGUCAC  CGCAGCCAGTG | -9.67 ± 0.59 | -81.63 ± 3.82 | -232.00 ± 10.92 | 49.4 | -9.34 ± 0.07 | -70.58 ± 0.64 | -197.47 ± 1.98 |
| CUAACAUGAGG  GATTGCACTCC | -7.31 ± 0.21 | -68.36 ± 1.78 | -196.85 ± 5.17 | 40.6 | -7.08 ± 0.05 | -62.01 ± 0.74 | -177.11 ± 2.35 |
| GUAGGAACAUG  CATCCCTGTAC | -8.54 ± 0.44 | -116.22 ± 4.38 | -347.18 ± 13.12 | 42.5 | -8.59 ± 0.001 | -116.25 ± 0.001 | -347.14 ± 0.002 |
| CAGUGACCAAG  GTCACCGGTTC | -9.81 ± 0.27 | -96.30 ± 2.37 | -278.87 ± 6.88 | 47.9 | -10.00 ± 0.01 | -95.85 ± 0.17 | -276.80 ± 0.52 |
| CCUAGAGCAUG  GGATCCCGTAC | -9.21 ± 0.11 | -100.24 ± 1.20 | -293.48 ± 3.54 | 45.5 | -9.37 ± 0.001 | -100.19 ± 0.029 | -292.81 ± 0.091 |
| CAGUGAUCAAG  GTCACCAGTTC | -8.11 ± 0.47 | -85.50 ± 3.57 | -249.52 ± 10.47 | 42.8 | -8.17 ± 0.02 | -83.02 ± 0.16 | -241.35 ± 0.52 |
| GUGUAAGUACC  CACACTCATGG | -6.81 ± 0.63 | -92.95 ± 4.63 | -277.72 ± 13.91 | 37.9 | -7.32 ± 0.01 | -92.71 ± 0.04 | -275.32 ± 0.14 |
| CAUGUACAGUG  GTACACGTCAC | -8.21 ± 0.49 | -99.08 ± 4.00 | -292.98 ± 11.86 | 42.3 | -8.90 ± 0.01 | -97.70 ± 0.11 | -286.31 ± 0.36 |
| GCACUAUACCG  CGTGACATGGC | -7.45 ± 0.18 | -98.39 ± 2.17 | -293.23 ± 6.49 | 39.9 | -7.83 ± 0.001 | -98.58 ± 0.014 | -292.60 ± 0.041 |
| **A/G Internal Mismatch** |  |  |  |  |  |  |  |
| GAGCAAACGUC  CTCGTGTGCAG | -9.24 ± 0.80 | -97.21 ± 5.62 | -283.64 ± 16.47 | 45.9 | -9.41 ± 0.01 | -97.76 ± 0.09 | -284.85 ± 0.298 |
| GUUGAACCUAC  CAACTGGGATG | -7.77 ± 0.32 | -72.99 ± 2.54 | -210.29 ± 7.38 | 42.4 | -7.61 ± 0.07 | -64.60 ± 0.81 | -183.77 ± 2.58 |
| GAUCAAUCCAC  CTAGTGAGGTG | -7.58 ± 0.33 | -87.65 ± 3.11 | -258.18 ± 9.20 | 40.8 | -7.88 ± 0.01 | -86.95 ± 0.19 | -254.96 ± 0.61 |
| CGACCAACUUG  GCTGGGTGAAC | -7.49 ± 0.44 | -72.27 ± 3.16 | -208.89 ± 9.21 | 41.2 | -7.60 ± 0.13 | -69.35 ± 1.21 | -199.10 ± 3.86 |
| CAUUCACUACG  GTAAGGGATGC | -6.36 ± 2.65 | -88.06 ± 9.86 | -263.41 ± 29.64 | 36.4 | -6.63 ± 0.01 | -88.07 ± 0.03 | -262.60 ± 0.11 |
| GCGUCAGUCAC  CGCAGGCAGTG | -9.73 ± 1.53 | -73.01 ± 6.47 | -204.01 ± 18.23 | 51.2 | -9.69 ± 0.23 | -72.67 ± 1.06 | -203.05 ± 3.29 |
| CUAACAUGAGG  GATTGGACTCC | -7.51 ± 0.30 | -75.59 ± 2.48 | -219.49 ± 7.26 | 41.1 | -7.49 ± 0.01 | -77.68 ± 0.12 | -226.28 ± 0.37 |
| GUAGGAACAUG  CATCCGTGTAC | -8.43 ± 0.38 | -78.30 ± 2.89 | -225.28 ± 8.37 | 44.7 | -8.17 ± 0.052 | -66.71 ± 0.56 | -188.75 ± 1.77 |
| CAGUGACCAAG  GTCACGGGTTC | -10.10 ± 0.75 | -84.66 ± 4.79 | -240.41 ± 13.67 | 50.7 | -9.81 ± 0.07 | -76.22 ± 0.60 | -214.12 ± 1.88 |
| CCUAGAGCAUG  GGATCGCGTAC | -9.01 ± 0.45 | -73.40 ± 3.17 | -207.63 ± 9.03 | 47.8 | -8.61 ± 0.06 | -63.35 ± 0.72 | -176.50 ± 2.25 |
| CAGUGAUCAAG  GTCACGAGTTC | -8.54 ± 0.14 | -93.69 ± 1.52 | -274.54 ± 4.49 | 43.8 | -8.76 ± 0.002 | -93.56 ± 0.05 | -273.42 ± 0.16 |
| GUGUAAGUACC  CACAGTCATGG | -6.88 ± 1.01 | -103.92 ± 6.41 | -312.87 ± 19.36 | 38.1 | -6.91 ± 0.002 | -103.92 ± 0.004 | -312.79 ± 0.01 |
| CAUGUACAGUG  GTACAGGTCAC | -8.16 ± 0.19 | -65.87 ± 1.48 | -186.07 ± 4.22 | 44.8 | -7.98 ± 0.03 | -56.17 ± 0.58 | -155.36 ± 1.81 |
| GCACUAUACCG  CGTGAGATGGC | -7.85 ± 0.90 | -81.11 ± 5.63 | -236.19 ± 16.48 | 42.1 | -7.65 ± 0.02 | -81.22 ± 0.11 | -237.23 ± 0.37 |
| **U/C Internal Mismatch** |  |  |  |  |  |  |  |
| GCACUAUACCG  CGTGATCTGGC | -7.41 ± 1.37 | -135.81 ± 9.47 | -413.98 ± 28.94 | 39.0 | -7.44 ± 0.001 | -135.83 ± 0.001 | -413.97 ± 0.001 |
| GAUCAAUCCAC  CTAGTTCGGTG | -7.39 ± 2.44 | -91.00 ± 10.51 | -269.60 ± 31.27 | 39.9 | -7.75 ± 0.04 | -89.60 ± 0.18 | -263.93 ± 0.56 |
| CUAACAUGAGG  GATTGTCCTCC | -7.52 ± 5.11 | -86.03 ± 13.77 | -253.13 ± 40.75 | 40.6 | -7.61 ± 0.03 | -84.72 ± 0.07 | -248.60 ± 0.24 |
| UAAGAUUGGUA  ATTCTCACCAT | -6.36 ± 1.55 | -82.82 ± 7.20 | -246.52 ± 21.57 | 36.4 | -6.55 ± 0.01 | -82.97 ± 0.05 | -246.40 ± 0.16 |
| UGGUCUGUAGA  ACCAGCCATCT | -7.87 ± 0.19 | -84.85 ± 1.91 | -248.21 ± 5.61 | 42.0 | -8.00 ± 0.005 | -84.53 ± 0.096 | -246.74 ± 0.307 |
| GAACCUCUCUA  CTTGGCGAGAT | -6.63 ± 0.49 | -99.72 ± 4.16 | -300.12 ± 12.58 | 37.3 | -6.77 ± 0.003 | -99.82 ± 0.007 | -300.00 ± 0.014 |
| GGAACUUCAUU  CCTTGCAGTAA | -6.67 ± 0.45 | -68.61 ± 3.45 | -199.74 ± 10.12 | 37.6 | -6.42 ± 0.05 | -61.25 ± 0.53 | -176.79 ± 1.71 |
| CAUGUACAGUG  GTACCTGTCAC | -8.20 ± 0.22 | -97.80 ± 2.28 | -288.89 ± 6.76 | 42.4 | -9.05 ± 0.01 | -95.52 ± 0.15 | -278.83 ± 0.49 |
| UACAGUCCAUG  ATGTCCGGTAC | -7.82 ± 0.39 | -79.75 ± 3.24 | -231.93 ± 9.49 | 42.1 | -7.72 ± 0.09 | -68.16 ± 1.09 | -194.87 ± 3.46 |
| AACGGUGCGAA  TTGCCCCGCTT | -10.05 ± 0.13 | -86.42 ± 1.12 | -246.24 ± 3.22 | 50.2 | -9.72 ± 0.02 | -74.93 ± 0.50 | -210.27 ± 1.55 |
| CUACGUUGCAA  GATGCCACGTT | -6.97 ± 0.08 | -95.19 ± 1.08 | -284.46 ± 3.25 | 38.4 | -7.51 ± 0.004 | -95.05 ± 0.125 | -282.25 ± 0.406 |
| UAAGAUUGGUA  ATTCTACCCAT | -6.33 ± 1.58 | -77.02 ± 7.72 | -227.92 ± 22.98 | 36.2 | -6.42 ± 0.02 | -77.11 ± 0.08 | -227.90 ± 0.27 |
| AGUCUUUCAGG  TCAGACAGTCC | -6.05 ± 0.49 | -94.01 ± 4.47 | -283.61 ± 13.53 | 35.4 | -6.39 ± 0.005 | -94.19 ± 0.012 | -283.11 ± 0.056 |
| **U/G Internal Mismatch** |  |  |  |  |  |  |  |
| GAUCAAUCCAC  CTAGTTGGGTG | -10.56 ± 0.11 | -102.32 ± 1.08 | -295.87 ± 3.13 | 49.7 | -10.54 ± 0.007 | -100.38 ± 0.185 | -289.66 ± 0.580 |
| CUAACAUGAGG  GATTGTGCTCC | -9.49 ± 0.49 | -94.54 ± 3.71 | -274.21 ± 10.83 | 47.0 | -9.65 ± 0.01 | -93.07 ± 0.15 | -268.98 ± 0.47 |
| UAAGAUUGGUA  ATTCTGACCAT | -7.78 ± 0.94 | -101.41 ± 5.99 | -301.90 ± 17.91 | 40.9 | -8.15 ± 0.002 | -101.64 ± 0.009 | -301.43 ± 0.023 |
| UGGUCUGUAGA  ACCAGGCATCT | -8.96 ± 0.29 | -74.29 ± 2.20 | -210.64 ± 6.30 | 47.5 | -8.71 ± 0.031 | -67.59 ± 0.403 | -189.86 ± 1.263 |
| GGAACUUCAUU  CCTTGGAGTAA | -7.86 ± 0.97 | -91.26 ± 6.26 | -268.91 ± 18.54 | 41.6 | -8.04 ± 0.006 | -91.54 ± 0.043 | -269.23 ± 0.140 |
| CAUGUACAGUG  GTACGTGTCAC | -9.55 ± 0.71 | -104.71 ± 5.00 | -306.82 ± 14.71 | 46.2 | -9.90 ± 0.004 | -104.91 ± 0.031 | -306.32 ± 0.091 |
| UACAGUCCAUG  ATGTCGGGTAC | -10.63 ± 0.24 | -102.03 ± 2.13 | -294.72 ± 6.18 | 50.0 | -10.87 ± 0.01 | -101.81 ± 0.15 | -293.21 ± 0.46 |
| AACGGUGCGAA  TTGCCGCGCTT | -12.40 ± 0.54 | -90.74 ± 3.59 | -252.60 ± 10.05 | 58.4 | -11.89 ± 0.03 | -80.84 ± 0.33 | -222.31 ± 1.02 |
| CUACGUUGCAA  GATGCGACGTT | -8.55 ± 0.31 | -66.35 ± 2.17 | -186.37 ± 6.17 | 46.7 | -8.44 ± 0.080 | -65.82 ± 0.960 | -185.02 ± 3.011 |
| CGCUUAACUAG  GCGAGTTGATC | -7.17 ± 0.23 | -127.77 ± 3.15 | -388.85 ± 9.60 | 38.6 | -7.86 ± 0.001 | -128.13 ± 0.001 | -387.77 ± 0.005 |
| UAAGAUUGGUA  ATTCTAGCCAT | -7.85 ± 0.32 | -89.05 ± 3.02 | -261.83 ± 8.92 | 41.7 | -8.07 ± 0.006 | -88.20 ± 0.086 | -258.35 ± 0.278 |
| GGAACUUCAUU  CCTTGAGGTAA | -8.03 ± 0.46 | -93.56 ± 3.73 | -275.78 ± 11.04 | 42.0 | -8.33 ± 0.004 | -93.64 ± 0.038 | -275.06 ± 0.118 |
| AGUCUUUCAGG  TCAGAGAGTCC | -6.98 ± 0.68 | -91.44 ± 4.79 | -272.31 ± 14.33 | 38.5 | -7.33 ± 0.004 | -91.69 ± 0.025 | -272.01 ± 0.086 |
| **C/A Internal Mismatch** |  |  |  |  |  |  |  |
| CUAACAUGAGG  GATTATACTCC | -6.61 ± 2.57 | -84.95 ± 10.49 | -252.57 ± 31.36 | 37.3 | -6.80 ± 0.042 | -82.54 ± 0.173 | -244.18 ± 0.561 |
| GUUGAACCUAC  CAACTTAGATG | -4.81 ± 0.21 | -141.87 ± 4.13 | -441.91 ± 12.88 | 33.3 | -4.93 ± 0.001 | -141.93 ± 0.001 | -441.71 ± 0.008 |
| AAUACCCACCG  TTATGAGTGGC | -7.10 ± 0.40 | -66.36 ± 3.05 | -191.06 ± 8.86 | 39.7 | -6.93 ± 0.08 | -56.56 ± 0.81 | -160.05 ± 2.59 |
| AAUGCCGUAUG  TTACGACATAC | -6.35 ± 0.96 | -81.48 ± 6.03 | -242.25 ± 18.04 | 36.3 | -6.57 ± 0.019 | -81.22 ± 0.127 | -240.70 ± 0.410 |
| CAUCGCAGCAA  GTAGCATCGTT | -8.06 ± 0.19 | -83.56 ± 1.84 | -243.44 ± 5.40 | 42.8 | -8.27 ± 0.01 | -84.22 ± 0.19 | -244.88 ± 0.61 |
| AAUGCCGUAUG  TTACAGCATAC | -6.87 ± 0.75 | -83.17 ± 5.24 | -246.00 ± 15.59 | 38.3 | -6.98 ± 0.008 | -82.91 ± 0.061 | -244.80 ± 0.196 |
| AAUAGCUCUGG  TTATCAAGACC | -6.33 ± 0.76 | -81.23 ± 5.28 | -241.48 ± 15.79 | 36.2 | -6.35 ± 0.04 | -75.02 ± 0.34 | -221.39 ± 1.10 |
| UACAGUCCAUG  ATGTCAAGTAC | -5.97 ± 0.92 | -93.73 ± 6.39 | -282.96 ± 19.36 | 35.2 | -6.24 ± 0.001 | -93.87 ± 0.007 | -282.56 ± 0.020 |
| ACGUUCGGAUC  TGCAAACCTAG | -8.09 ± 0.05 | -89.74 ± 0.56 | -263.28 ± 1.65 | 42.5 | -8.23 ± 0.003 | -89.65 ± 0.081 | -262.50 ± 0.252 |
| UGGUCUGUAGA  ACCAAACATCT | -6.41 ± 0.17 | -82.50 ± 1.87 | -245.33 ± 5.60 | 36.5 | -6.60 ± 0.003 | -82.55 ± 0.054 | -244.87 ± 0.173 |
| **C/T Internal Mismatch** |  |  |  |  |  |  |  |
| CUAACAUGAGG  GATTTTACTCC | -5.99 ± 0.98 | -97.81 ± 6.74 | -296.02 ± 20.48 | 35.3 | -6.13 ± 0.001 | -97.89 ± 0.005 | -295.86 ± 0.015 |
| GUUGAACCUAC  CAACTTTGATG | -5.44 ± 0.99 | -114.23 ± 7.39 | -350.77 ± 22.75 | 34.1 | -5.49 ± 0.010 | -114.26 ± 0.001 | -350.70 ± 0.035 |
| AAUACCCACCG  TTATGGTTGGC | -6.80 ± 0.27 | -76.89 ± 2.61 | -225.99 ± 7.73 | 38.1 | -7.00 ± 0.12 | -66.74 ± 1.75 | -192.64 ± 5.60 |
| AAUACCCACCG  TTATGTGTGGC | -7.23 ± 0.34 | -79.18 ± 3.02 | -232.00 ± 8.90 | 39.8 | -7.22 ± 0.01 | -78.18 ± 0.13 | -228.78 ± 0.40 |
| AAUGCCGUAUG  TTACGTCATAC | -6.37 ± 0.15 | -104.10 ± 2.11 | -315.08 ± 6.41 | 36.5 | -6.39 ± 0.002 | -104.10 ± 0.001 | -315.04 ± 0.009 |
| CAUCGCAGCAA  GTAGCTTCGTT | -7.64 ± 0.32 | -95.94 ± 3.02 | -284.73 ± 8.99 | 40.6 | -8.12 ± 0.002 | -96.15 ± 0.027 | -283.82 ± 0.083 |
| AAUGCCGUAUG  TTACTGCATAC | -6.80 ± 0.19 | -84.64 ± 2.11 | -250.96 ± 6.30 | 38.0 | -6.99 ± 0.002 | -84.72 ± 0.042 | -250.61 ± 0.132 |
| AAUAGCUCUGG  TTATCTAGACC | -5.17 ± 0.71 | -113.27 ± 6.18 | -348.55 ± 19.08 | 33.3 | -5.53 ± 0.001 | -113.52 ± 0.002 | -348.17 ± 0.007 |
| UACAGUCCAUG  ATGTCATGTAC | -6.39 ± 0.28 | -79.69 ± 2.78 | -236.32 ± 8.30 | 36.5 | -6.78 ± 0.011 | -79.65 ± 0.153 | -234.95 ± 0.493 |
| ACGUUCGGAUC  TGCAATCCTAG | -7.83 ± 0.35 | -101.33 ± 3.53 | -301.49 ± 10.54 | 41.0 | -8.34 ± 0.002 | -101.60 ± 0.020 | -300.69 ± 0.060 |
| UGGUCUGUAGA  ACCATACATCT | -6.74 ± 0.85 | -80.41 ± 5.54 | -237.54 ± 16.45 | 37.8 | -6.81 ± 0.008 | -79.68 ± 0.060 | -234.97 ± 0.194 |
| **G/A Internal Mismatch** |  |  |  |  |  |  |  |
| CCUAGAGCAUG  GGATATCGTAC | -8.13 ± 0.66 | -78.14 ± 4.17 | -225.71 ± 12.12 | 43.5 | -8.02 ± 0.107 | -66.60 ± 0.822 | -188.88 ± 2.594 |
| CCUAGAGCAUG  GGATCTAGTAC | -8.16 ± 0.63 | -69.92 ± 3.78 | -199.15 ± 10.87 | 44.4 | -7.89 ± 0.049 | -62.45 ± 0.372 | -175.91 ± 1.172 |
| UUCCAGGAAGG  AAGGTACTTCC | -10.36 ± 0.51 | -114.97 ± 4.16 | -337.28 ± 12.24 | 47.7 | -11.07 ± 0.004 | -115.38 ± 0.037 | -336.30 ± 0.115 |
| GUGUAAGUACC  CACATTAATGG | -7.69 ± 0.30 | -115.79 ± 3.44 | -348.53 ± 10.38 | 40.1 | -7.69 ± 0.008 | -115.76 ± 0.005 | -348.45 ± 0.041 |
| CAUCGCAGCAA  GTAGAGTCGTT | -8.24 ± 0.29 | -71.32 ± 2.35 | -203.41 ± 6.76 | 44.6 | -8.03 ± 0.05 | -61.31 ± 0.742 | -171.78 ± 2.335 |
| ACGUUCGGAUC  TGCAAGACTAG | -7.66 ± 0.04 | -63.40 ± 0.38 | -179.70 ± 1.09 | 42.6 | -7.26 ± 0.01 | -54.08 ± 0.62 | -150.96 ± 1.98 |
| GUAGGAACAUG  CATCATTGTAC | -7.41 ± 0.61 | -102.99 ± 4.74 | -308.17 ± 14.24 | 39.7 | -7.42 ± 0.001 | -102.98 ± 0.003 | -308.11 ± 0.008 |
| AACGGUGCGAA  TTGCAACGCTT | -9.66 ± 0.54 | -101.49 ± 4.11 | -296.07 ± 12.03 | 46.9 | -10.00 ± 0.007 | -101.30 ± 0.071 | -294.39 ± 0.224 |
| CAGUGACCAAG  GTCAATGGTTC | -7.55 ± 0.50 | -104.73 ± 4.25 | -313.34 ± 12.75 | 40.0 | -7.64 ± 0.001 | -104.79 ± 0.003 | -313.23 ± 0.006 |
| AACGGUGCGAA  TTGCCAAGCTT | -9.00 ± 0.10 | -95.41 ± 1.11 | -278.61 ± 3.27 | 45.2 | -9.15 ± 0.002 | -94.59 ± 0.05 | -275.47 ± 0.17 |
| UGGUCUGUAGA  ACCAGAAATCT | -5.82 ± 0.08 | -100.72 ± 1.28 | -305.99 ± 3.90 | 34.8 | -6.02 ± 0.001 | -100.91 ± 0.004 | -305.96 ± 0.009 |
| **G/T Internal Mismatch** |  |  |  |  |  |  |  |
| CCUAGAGCAUG  GGATTTCGTAC | -9.87 ± 0.30 | -96.94 ± 2.62 | -280.73 ± 7.62 | 48.1 | -9.98 ± 0.01 | -95.72 ± 0.14 | -276.45 ± 0.43 |
| CCUAGAGCAUG  GGATCTTGTAC | -10.21 ± 0.32 | -91.39 ± 2.55 | -261.77 ± 7.35 | 50.0 | -9.78 ± 0.03 | -78.55 ± 0.40 | -221.72 ± 1.25 |
| UUCCAGGAAGG  AAGGTTCTTCC | -10.29 ± 0.15 | -82.80 ± 1.22 | -233.80 ± 3.46 | 51.7 | -9.87 ± 0.02 | -70.82 ± 0.48 | -196.52 ± 1.49 |
| GUGUAAGUACC  CACATTTATGG | -8.27 ± 0.11 | -80.98 ± 1.05 | -234.43 ± 3.07 | 43.8 | -8.03 ± 0.02 | -69.53 ± 0.55 | -198.30 ± 1.74 |
| UCACACGAUAA  AGTGTGTTATT | -8.21 ± 0.93 | -96.24 ± 6.27 | -283.82 ± 18.58 | 42.5 | -8.55 ± 0.006 | -96.30 ± 0.045 | -282.93 ± 0.142 |
| CAUCGCAGCAA  GTAGTGTCGTT | -10.68 ± 0.26 | -79.84 ± 1.86 | -222.99 ± 5.22 | 54.0 | -10.35 ± 0.02 | -73.34 ± 0.24 | -203.09 ± 0.75 |
| ACGUUCGGAUC  TGCAAGTCTAG | -9.97 ± 0.42 | -90.94 ± 3.19 | -261.06 ± 9.19 | 49.2 | -9.59 ± 0.04 | -77.34 ± 0.42 | -218.44 ± 1.30 |
| CUACGUUGCAA  GATGTAACGTT | -9.12 ± 0.32 | -95.25 ± 2.95 | -277.69 ± 8.65 | 45.7 | -9.27 ± 0.005 | -95.78 ± 0.08 | -278.93 ± 0.25 |
| GUAGGAACAUG  CATCTTTGTAC | -9.37 ± 0.63 | -102.20 ± 4.59 | -299.29 ± 13.50 | 45.9 | -9.70 ± 0.003 | -102.52 ± 0.027 | -299.28 ± 0.086 |
| AACGGUGCGAA  TTGCTACGCTT | -11.82 ± 0.20 | -88.18 ± 1.50 | -246.21 ± 4.22 | 56.8 | -11.35 ± 0.02 | -78.01 ± 0.25 | -214.94 ± 0.77 |
| CAGUGACCAAG  GTCATTGGTTC | -10.00 ± 0.81 | -86.09 ± 5.15 | -245.34 ± 14.77 | 50.0 | -9.66 ± 0.04 | -73.53 ± 0.35 | -205.94 ± 1.09 |
| AACGGUGCGAA  TTGCCATGCTT | -11.48 ± 0.23 | -85.88 ± 1.69 | -239.88 ± 4.75 | 56.0 | -11.15 ± 0.02 | -78.52 ± 0.30 | -217.22 ± 0.90 |
| UGGUCUGUAGA  ACCAGATATCT | -9.22 ± 0.41 | -97.59 ± 3.60 | -284.92 ± 10.55 | 45.8 | -9.45 ± 0.006 | -97.53 ± 0.083 | -284.00 ± 0.266 |
