## Supplementary Table S2 for "Thermodynamic Parameters Contributions of Single Internal Mismatches In RNA/DNA Hybrid Duplexes"

| **Table S2:** Thermodynamics of duplex formation of RNA/DNA Hybrid Duplexes containing internal mismatches. All Tm’s are calculated using a duplex concentration of 1e^-4^M. | | | | | | | |
| --- | --- | --- | --- | --- | --- | --- | --- |
| **RNA/DNA Duplex** | **1/Tm Method Curve fit Method** | | | | | | |
| **Control Sequences** | **ΔG°_37_ (kcal/mol)** | **ΔH° (kcal/mol)** | **ΔS° (cal/K**mol)** | **T_m_ (°C)** | **ΔG°_37_ (kcal/mol)** | **ΔH° (kcal/mol)** | **ΔS° (cal/K**mol)** |
| AGGUAGGU*  TCCATCCA | -8.67 ± 0.06 | -68.6 ± 4.5 | -193.1 ± 14.3 | 47.2 | -8.67 ± 0.07 | -69.4 ± 3.4 | -195.8 ± 10.8 |
| GCUCAACCCG*  CGAGTTGGGC | -11.16 ± 0.47 | -68.0 ± 8.4 | -183.2 ± 25.9 | 59.6 | -11.61 ± 0.16 | -77.4 ± 2.4 | -212.0 ± 7.4 |
| GGCGAUGAUG*  CCGCTACTAC | -12.28 ± 0.71 | -90.42 ± 13.0 | -252.0 ± 8.1 | 58.0 | -12.10 ± 0.12 | -86.8 ± 2.6 | -240.7 ± 8.1 |
| CGAGGAUGGC*  GCTCCTACCG | -13.89 ± 0.55 | -83.9 ± 7.9 | -225.8 ± 23.6 | 66.8 | -14.46 ± 0.24 | -92.2 ± 2.6 | -250.5 ± 7.6 |
| GUCCUGCUCA*  CAGGACGAGT | -10.92 ± 1.31 | -77.3 ± 21.7 | -213.9 ± 66.8 | 55.7 | -10.75 ± 0.22 | -73.1 ± 4.0 | -200.9 ± 12.2 |
| CCACACAGAG*  GGTGTGTCTC | -11.65 ± 0.42 | -80.8 ± 8.3 | -222.9 ± 25.5 | 58.0 | -11.81 ± 0.16 | -84.1 ± 3.1 | -232.9 ± 9.5 |
| CAUGAGCUAC*  GTACTCGATG | -10.17 ± 0.11 | -63.9 ± 3.0 | -173.2 ± 9.3 | 55.7 | -10.67 ± 0.12 | -78.8 ± 2.2 | -219.7 ± 7.0 |
| GAGACACACC*  CTCTGTGTGG | -12.41 ± 0.85 | -88.0 ± 14.5 | -243.6 ± 44.1 | 59.2 | -12.36 ± 0.14 | -86.9 ± 2.1 | -240.4 ± 6.3 |
| GAACUCUGUG*  CTTGAGACAC | -8.42 ± 0.03 | -65.7 ± 2.7 | -184.6 ± 8.6 | 46.2 | -8.44 ± 0.05 | -67.9 ± 1.3 | -191.8 ± 4.1 |
| GGACGACG*  CCTGCTGC | -10.11 ± 0.19 | -63.9 ± 4.7 | -173.3 ± 14.5 | 55.4 | -10.32 ± 0.08 | -69.6 ± 1.6 | -191.0 ± 5.0 |
| CGCAGUCCAC*  GCGTCAGGTG | -13.72 ± 0.55 | -92.4 ± 8.4 | -253.7 ± 25.2 | 63.2 | -13.77 ± 0.12 | -93.3 ± 1.6 | -256.3 ± 4.9 |
| **A/A Internal Mismatch** |  |  |  |  |  |  |  |
| CCACAACAGAG*  GGTGTAGTCTC | -8.23 ± 0.02 | -74.6 ± 2.3 | -214.0 ± 7.4 | 44.2 | -8.27 ± 0.07 | -84.9 ± 5.3 | -247.2 ± 16.8 |
| GUCCUAGCUCA*  CAGGAACGAGT | -7.38 ± 0.07 | -74.5 ± 4.5 | -216.5 ± 14.7 | 40.6 | -7.40 ± 0.06 | -71.7 ± 1.9 | -207.4 ± 6.1 |
| CAUGAAGCUAC*  GTACTACGATG | -7.64 ± 0.09 | -79.0 ± 5.6 | -230.0 ± 18.3 | 41.4 | -7.60 ± 0.06 | -86.7 ± 3.6 | -255.1 ± 11.5 |
| GAGACAACACC*  CTCTGATGTGG | -8.49 ± 0.35 | -80.0 ± 11.8 | -230.6 ± 37.7 | 44.8 | -8.46 ± 0.12 | -74.7 ± 5.8 | -213.5 ± 18.8 |
| CGAGGAAUGGC*  GCTCCATACCG | -9.88 ± 0.29 | -86.6 ± 9.5 | -247.3 ± 29.8 | 49.5 | -9.85 ± 0.09 | -85.0 ± 2.5 | -242.3 ± 7.9 |
| GAACUACUGUG*  CTTGAAGACAC | -6.03 ± 0.17 | -62.5 ± 5.0 | -182.1 ± 16.7 | 34.5 | -5.83 ± 0.03 | -69.0 ± 2.1 | -203.7 ± 6.7 |
| AGGUAAGGU*  TCCAATCCA | -5.46 ± 0.01 | -59.9 ± 0.2 | -175.6 ± 0.5 | 31.5 | -5.35 ± 0.14 | -62.8 ± 3.0 | -185.1 ± 10.3 |
| GCUCAAACCCG*  CGAGTATGGGC | -8.40 ± 0.05 | -66.7 ± 3.8 | -188.1 ± 12.3 | 45.8 | -8.48 ± 0.10 | -75.3 ± 6.8 | -215.4 ± 21.8 |
| GCCGUAUCAAC*  CGGCAAAGTTG | -7.72 ± 0.17 | -65.9 ± 7.2 | -187.5 ± 23.5 | 42.7 | -7.67 ± 0.11 | -75.7 ± 1.1 | -219.4 ± 3.8 |
| CGCAGAUCCAC*  GCGTCAAGGTG | -8.87 ± 0.13 | -70.0 ± 6.7 | -197.2 ± 21.2 | 47.6 | -8.97 ± 0.08 | -77.7 ± 4.1 | -221.5 ± 13.1 |
| AAGCAGUAG**  TTCGACATC | -4.95 ± 0.09 | -58.1 ± 1.3 | -171.2 ± 4.0 | 29.0 | -5.10 ± 0.17 | -54.8 ± 1.9 | -160.2 ± 6.7 |
| UCACACUAG**  AGTGAGATC | -3.99 ± 0.32 | -42.6 ± 2.3 | -124.5 ± 6.9 | 19.5 | -3.71 ± 0.38 | -44.5 ± 3.0 | -131.4 ± 9.8 |
| UGAGAGUAC**  ACTCACATG | -5.92 ± 0.20 | -67.4 ± 2.0 | -198.1 ± 6.1 | 34.4 | -5.87 ± 0.37 | -69.6 ± 8.8 | -205.4 ± 28.6 |
| UUGGACACC**  AACCAGTGG | -5.71 ± 0.41 | -45.3 ± 2.6 | -127.6 ± 7.5 | 31.6 | -5.62 ± 0.11 | -48.4 ± 2.5 | -137.8 ± 8.2 |
| **C/C Internal Mismatch** |  |  |  |  |  |  |  |
| GGACCGACG*  CCTGCCTGC | -6.88 ± 0.88 | -51.0 ± 3.1 | -142.1 ± 10.2 | 39.2 | -6.80 ± 0.07 | -55.3 ± 2.5 | -156.4 ± 8.1 |
| CCACACCAGAG*  GGTGTCGTCTC | -7.25 ± 0.07 | -65.7 ± 3.9 | -188.6 ± 12.7 | 40.2 | -7.26 ± 0.10 | -67.3 ± 5.9 | -193.7 ± 19.3 |
| GUCCUCGCUCA*  CAGGACCGAGT | -6.84 ± 0.20 | -71.8 ± 7.5 | -209.4 ± 24.4 | 38.3 | -6.79 ± 0.08 | -75.7 ± 5.7 | -222.1 ± 18.3 |
| CAUGACGCUAC*  GTACTCCGATG | -6.88 ± 0.26 | -79.1 ± 11.6 | -232.9 ± 37.7 | 38.4 | -6.79 ± 0.04 | -86.4 ± 1.0 | -256.7 ± 3.1 |
| GAGACCACACC*  CTCTGCTGTGG | -7.83 ± 0.20 | -62.7 ± 7.0 | -177.0 ± 22.7 | 43.5 | -7.80 ± 0.11 | -71.5 ± 1.7 | -205.4 ± 5.3 |
| CGAGGCAUGGC*  GCTCCCTACCG | -9.75 ± 0.14 | -80.1 ± 5.4 | -226.9 ± 17.1 | 50.0 | -9.82 ± 0.04 | -84.0 ± 4.6 | -239.0 ± 14.7 |
| AGGUCAGGU*  TCCACTCCA | -5.49 ± 1.11 | -53.8 ± 16.7 | -155.6 ± 55.5 | 31.4 | -5.24 ± 0.07 | -59.8 ± 4.3 | -175.8 ± 14.0 |
| GGCGACUGAUG*  CCGCTCACTAC | -7.74 ± 0.04 | -76.8 ± 3.8 | -222.5 ± 12.2 | 41.9 | -7.70 ± 0.06 | -88.8 ± 3.0 | -261.5 ± 9.7 |
| GACUGCCUCCA*  CTGACCGAGGT | -7.55 ± 0.19 | -60.7 ± 7.6 | -171.4 ± 24.7 | 42.2 | -7.59 ± 0.15 | -61.0 ± 8.3 | -172.2 ± 27.1 |
| CGAGCGAUGGC*  GCTCCCTACCG | -9.87 ± 0.38 | -88.0 ± 12.5 | -252.0 ± 39.4 | 49.1 | -9.88 ± 0.13 | -88.9 ± 5.9 | -254.7 ± 18.6 |
| CGCAGCUCCAC*  GCGTCCAGGTG | -8.57 ± 0.59 | -81.1 ± 18.2 | -233.7 ± 58.0 | 45.2 | -8.65 ± 0.13 | -89.9 ± 7.2 | -261.9 ± 22.9 |
| **G/G Internal Mismatch** |  |  |  |  |  |  |  |
| CCACAGCAGAG*  GGTGTGGTCTC | -9.96 ± 0.40 | -70.2 ± 10.1 | -194.3 ± 31.4 | 52.9 | -10.30 ± 0.16 | -81.3 ± 3.8 | -228.9 ± 11.7 |
| GUCCUGGCUCA*  CAGGAGCGAGT | -8.32 ± 0.22 | -66.1 ± 8.3 | -186.1 ± 26.5 | 45.6 | -8.37 ± 0.08 | -73.4 ± 8.4 | -209.5 ± 26.9 |
| CAUGAGGCUAC*  GTACTGCGATG | -9.00 ± 0.10 | -76.1 ± 5.1 | -216.2 ± 16.3 | 47.4 | -9.13 ± 0.07 | -87.5 ± 2.2 | -252.6 ± 7.3 |
| GAGACGACACC*  CTCTGGTGTGG | -9.81 ± 0.16 | -82.2 ± 6.0 | -233.4 ± 18.8 | 49.9 | -9.91 ± 0.07 | -87.1 ± 3.0 | -248.7 ± 9.7 |
| CGAGGGAUGGC*  GCTCCGTACCG | -12.65 ± 0.61 | -92.9 ± 11.3 | -258.6 ± 34.6 | 58.9 | -12.65 ± 0.24 | -93.0 ± 4.7 | -259.1 ± 14.4 |
| GAACUGCUGUG*  CTTGAGGACAC | -7.44 ± 0.32 | -73.0 ± 11.2 | -211.5 ± 36.3 | 40.8 | -7.41 ± 0.08 | -77.6 ± 2.9 | -226.4 ± 9.3 |
| AGGUGAGGU*  TCCAGTCCA | -7.13 ± 0.06 | -60.2 ±3.8 | -171.2 ± 12.5 | 40.2 | -7.03 ± 0.06 | -67.8 ± 1.8 | -196.0 ± 5.9 |
| GGCGAGUGAUG*  CCGCTGACTAC | -9.60 ± 0.09 | -76.5 ± 3.4 | -215.8 ± 10.9 | 50.0 | -9.87 ± 0.07 | -89.5 ± 3.1 | -256.6 ± 9.8 |
| GCCACGUCACG*  CGGTGGAGTGC | -10.03 ± 0.08 | -89.0 ± 3.1 | -254.6 ±9.8 | 49.7 | -10.30 ± 0.14 | -99.8 ± 5.8 | -288.7 ± 18.1 |
| CGCAGGUCCAC*  GCGTCGAGGTG | -10.96 ± 0.18 | -88.0 ± 5.2 | -248.4 ± 16.2 | 53.4 | -11.31 ± 0.38 | -98.8 ± 12.4 | -281.9 ± 38.8 |
| AAGCGGUAG**  TTCGGCATC | -5.89 ± 0.54 | -73.6 ± 3.9 | -218.2 ± 11.7 | 34.5 | -5.81 ± 0.20 | -72.5 ± 2.5 | -214.9 ± 8.0 |
| UCACGCUAG**  AGTGGGATC | -5.28 ± 0.17 | -64.2 ± 1.9 | -190.0 ± 6.0 | 31.0 | -5.20 ± 0.09 | -67.0 ± 2.7 | -199.2 ± 8.7 |
| UGAGGGUAC**  ACTCGCATG | -7.38 ± 0.68 | -85.1 ± 4.6 | -250.7 ± 14.0 | 40.0 | -7.40 ± 0.11 | -85.5 ± 4.5 | -252.0 ± 14.8 |
| UUGGGCACC**  AACCGGTGG | -8.25 ± 0.40 | -61.9 ± 2.7 | -173.1 ± 7.8 | 45.7 | -8.26 ± 0.10 | -58.2 ± 3.6 | -161.1 ± 11.9 |
| **U/T Internal Mismatch** |  |  |  |  |  |  |  |
| GGACUGACG*  CCTGTCTGC | -7.36 ± 0.02 | -66.5 ± 1.9 | -190.7 ± 6.0 | 40.9 | -7.34 ± 0.04 | -72.8 ± 5.8 | -211.0 ± 18.7 |
| CCACAUCAGAG*  GGTGTTGTCTC | -8.83 ± 0.03 | -72.0 ± 2.0 | -203.6 ± 6.3 | 47.3 | -8.93 ± 0.05 | -81.2 ± 2.4 | -233.0 ± 7.8 |
| GUCCUUGCUCA*  CAGGATCGAGT | -7.84 ± 0.12 | -75.2 ± 6.9 | -217.2 ± 22.2 | 42.5 | -7.81 ± 0.07 | -81.4 ± 3.8 | -237.3 ± 12.1 |
| CAUGAUGCUAC*  GTACTTCGATG | -8.19 ± 0.03 | -85.3 ± 3.9 | -248.5 ± 12.6 | 43.1 | -8.22 ± 0.07 | -93.1 ± 5.2 | -273.8 ± 16.8 |
| GAGACUACACC*  CTCTGTTGTGG | -8.74 ± 0.16 | -72.8± 6.7 | -206.6 ± 21.4 | 46.7 | -8.86 ± 0.11 | -83.6 ± 3.5 | -240.9 ± 11.1 |
| CGAGGUAUGGC*  GCTCCTTACCG | -12.21 ± 1.41 | -107.4 ± 27.4 | -306.9 ± 84.4 | 54.3 | -12.07 ± 0.08 | -104.0 ± 2.3 | -296.3 ± 7.2 |
| GAACUUCUGUG*  CTTGATGACAC | -6.23 ± 0.14 | -67.3 ± 4.8 | -197.0 ± 16.0 | 35.6 | -6.13 ± 0.10 | -70.9 ± 4.7 | -208.9 ± 15.5 |
| AGGUUAGGU*  TCCATTCCA | -5.83 ± 0.18 | -62.6 ± 5.5 | -183.0 ± 18.3 | 33.6 | -5.58 ± 0.11 | -71.8 ± 3.7 | -213.5 ± 12.1 |
| GCUCAUACCCG*  CGAGTTTGGGC | -8.68 ± 0.31 | -76.5 ± 11.9 | -218.7 ± 37.9 | 45.9 | -8.81 ± 0.09 | -86.4 ± 4.3 | -250.2 ± 13.7 |
| GCCGUUUCAAC*  CGGCATAGTTG | -8.66 ± 0.25 | -81.8 ± 11.5 | -235.9 ± 36.5 | 45.2 | -8.72 ± 0.06 | -87.3 ± 2.5 | -253.3 ± 7.9 |
| GCCACUUCACG*  CGGTGTAGTGC | -9.65 ± 0.12 | -81.1 ± 5.2 | -230.3 ± 16.6 | 49.5 | -9.83 ± 0.10 | -90.2 ± 2.3 | -259.3 ± 7.3 |
| CGCAGUUCCAC*  GCGTCTAGGTG | -10.15 ± 0.25 | -99.9± 9.3 | -289.2 ± 29.3 | 48.7 | -10.11 ± 0.07 | -98.0 ± 2.3 | -283.4 ± 7.3 |
| UCACUCUAG**  AGTGTGATC | -4.06 ± 0.28 | -49.5 ± 2.1 | -146.5 ± 6.3 | 22.3 | -4.27 ± 0.22 | -47.0 ± 3.1 | -137.6 ± 10.7 |
| UGAGUGUAC**  ACTCTCATG | -5.68 ± 0.40 | -82.6 ± 3.5 | -248.0 ± 10.8 | 33.8 | -5.78 ± 0.21 | -78.6 ± 7.1 | -234.7 ± 23.6 |
| UUGGUCACC**  AACCTGTGG | -6.22 ± 0.11 | -56.6 ± 1.3 | -162.6 ± 3.8 | 35.0 | -6.06 ± 0.26 | -61.1 ± 3.3 | -174.4 ± 10.9 |
| *Watkins, N. E., Kennelly, W. J., Tsay, M. J., Tuin, A., Swenson, et al. (2011) Thermodynamic contributions of single internal rA·dA, rC·dC, rG·dG and rU·dT mismatches in RNA/DNA duplexes. *Nucleic Acids Research*, **39(5)**, 1894–1902.  **Sugimoto, N., Nakano, M. and Nakano, S. (2000) Thermodynamics-structure relationship of single mismatches in RNA/DNA duplexes. *Biochemistry*, **39**, 11270-11281. | | | | | | | |
