## Supplementary Table S3 for "Thermodynamic Parameters Contributions of Single Internal Mismatches In RNA/DNA Hybrid Duplexes"

| **Table S3:** Thermodynamics of duplex formation of paired RNA/DNA Hybrid Duplexes. All Tm’s are calculated using a duplex concentration of 1e^-4^M. | | | | |
| --- | --- | --- | --- | --- |
| **RNA sequence** | **ΔG°_37_ (kcal/mol)** | **ΔH° (kcal/mol)** | **ΔS° (cal/K·mol)** | **T_m_ (°C)** |
| AGCCG | -5.7 | -41.6 | -116 | 30.8 |
| CGGCU | -4.9 | -45.8 | -132 | 26.1 |
| GGUGG | -5.2 | -46 | -132 | 28 |
| ACCGCA | -6.4 | -45.4 | -126 | 36.1 |
| CAAUCG | -3.3 | -46.7 | -140 | 16.9 |
| CACGGC | -6.9 | -48.9 | -135 | 39.5 |
| CGAUUG | -2.6 | -49.3 | -151 | 14.1 |
| CGGUGC | -7 | -48.9 | -135 | 40.1 |
| CGUGCC | -6.3 | -49.3 | -139 | 35.3 |
| GCACCG | -6.2 | -49.7 | -141 | 34.8 |
| GGCACG | -7.3 | -51.8 | -144 | 41.5 |
| AAUACCG | -4.7 | -54.1 | -160 | 26.6 |
| ACGUAUG | -5 | -58.3 | -172 | 29.6 |
| AGCUUCA | -4.7 | -55.6 | -164 | 26.2 |
| CACGGCU | -8.5 | -53.9 | -147 | 48.6 |
| CAUACGU | -4.3 | -53.2 | -158 | 24.5 |
| GGACUUA | -4.9 | -43.8 | -125 | 25.9 |
| UAAGUCC | -5.4 | -46.2 | -132 | 29.4 |
| UGAAGCU | -6.4 | -43 | -118 | 36.5 |
| AAAAAAAA | -3.8 | -54 | -162 | 22.1 |
| AAGCGUAG | -7.8 | -67.2 | -192 | 43.1 |
| AAUCCAGU | -6.1 | -55.9 | -161 | 34.5 |
| AAUGUCGC | -7.2 | -64.9 | -186 | 39.9 |
| ACCUAGUC | -6.9 | -56 | -159 | 39.2 |
| ACGACCUC | -8.6 | -55 | -150 | 48.9 |
| ACUGGAUU | -7.2 | -57.8 | -163 | 40.9 |
| AGCGUAAG | -7.3 | -60.1 | -170 | 41.1 |
| AGUCCUGA | -6.7 | -54.7 | -155 | 37.9 |
| CAACAGCA | -7.8 | -52.7 | -145 | 44.8 |
| CACGGCUC | -9.6 | -71.6 | -200 | 50.5 |
| CGCUGUAA | -7.2 | -60.4 | -172 | 40.2 |
| CUACGCUU | -6.8 | -54.3 | -153 | 38.9 |
| CUAGUGGA | -8.3 | -63.4 | -178 | 45.1 |
| CUCACGGC | -9.6 | -70.3 | -196 | 51 |
| CUGAGUCC | -7.8 | -60.8 | -171 | 43.4 |
| CUUACGCU | -6.4 | -52.3 | -148 | 36.2 |
| GACUAGGU | -8.1 | -57 | -158 | 45.9 |
| GAGCCGUG | -9.5 | -67.3 | -187 | 51.4 |
| GAGGUCGU | -9.2 | -71.9 | -202 | 48.9 |
| GCCAGUUA | -7.2 | -62.9 | -180 | 40 |
| GCCGUGAG | -9.7 | -71.4 | -199 | 51.4 |
| GCGACAUU | -7.9 | -60.6 | -170 | 43.9 |
| UAACUGGC | -8.6 | -62.6 | -174 | 47.7 |
| UCCACUAG | -6.9 | -64.1 | -184 | 38.6 |
| UGCUGUUG | -6.7 | -52 | -146 | 38.3 |
| UGUUCGAC | -7.4 | -66.2 | -190 | 41.3 |
| UUACAGCG | -7 | -58.4 | -166 | 39.5 |
| UUGGCACC | -8.7 | -56.2 | -153 | 49.8 |
| AUAACUGGC | -8.7 | -60.7 | -168 | 48.7 |
| AUCUAUCCG | -6.8 | -59.9 | -171 | 38.4 |
| CAACAGCAA | -8.7 | -63.3 | -177 | 47.8 |
| CAACAGCAU | -9 | -71 | -200 | 48.1 |
| CGCUGUUAC | -8.2 | -71.5 | -205 | 44.2 |
| CGCUGUUAG | -8 | -67.7 | -193 | 44 |
| CUAACAGCG | -9 | -70.8 | -199 | 48.1 |
| GCCAGUUAA | -7.7 | -63.1 | -179 | 42.7 |
| GUAACAGCG | -9.1 | -77.5 | -221 | 47.8 |
| UUAACUGGC | -8.9 | -67.3 | -189 | 48.3 |
| ACGUAUUAUGC | -9.1 | -95.4 | -279 | 45.6 |
| GCAUAAUACGU | -10 | -97.3 | -282 | 48.4 |
| AAUGGAUUACAA | -10.2 | -90.7 | -260 | 50.2 |
| AUUGGAUACAAA | -10.8 | -93.6 | -267 | 51.9 |
| GUCAGGAAUCUG | -11.4 | -79.8 | -220 | 57.4 |
| UUGUAAUCCAUU | -8.5 | -76.9 | -221 | 45.4 |
| CGUGC | -2.3 | -48.3 | -149 | 14.2 |
| GCACG | -5 | -46.8 | -134 | 27.1 |
| UUUGUAUCCAAU | -8.8 | -84.9 | -246 | 45.6 |
| AGCCUAACUCAGC | -12.1 | -85.2 | -236 | 59.3 |
| Frank, J., Blocker, R., Marky, H., Freier, L. A., Kierzek, S. M., Jaeger, R., and Sugimoto, J. A., (1995), Thermodynamic Parameters to Predict Stability of RNA/DNA Hybrid Duplexes, *Proc. Natl. Acad. Sci.*, **34**, 3746-3750 | | | | |
