## Supplementary figures and images for "Thermodynamic Parameters Contributions of Single Internal Mismatches In RNA/DNA Hybrid Duplexes"

### Supplementary Figure S1

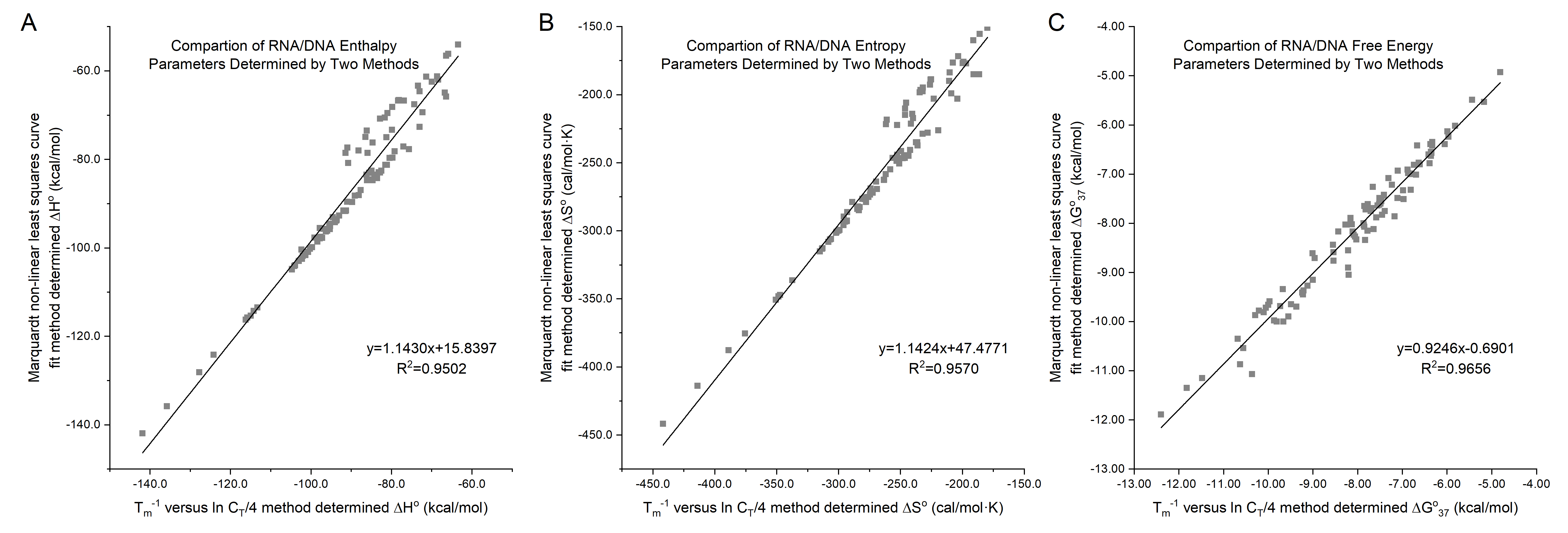
